# iPLA_2_-VIA is required for healthy aging in neurons, muscle, and female germline in *Drosophila melanogaster*

**DOI:** 10.1101/2021.04.25.441319

**Authors:** Surya Banerjee, Adina Schonbrun, Sogol Eizadshenass, Shimshon Benji, Yaakov Tzvi Cantor, Liam Eliach, Matthew Lubin, Zev Narrowe, Jeremy Purow, Benjamin Shulman, Leib Wiener, Josefa Steinhauer

## Abstract

Neurodegenerative disease (ND) is a growing health burden worldwide, but its causes and treatments remain elusive. Although most cases of ND are sporadic, rare familial cases have been attributed to single genes, which can be investigated in animal models. We have generated a new mutation in the calcium-independent phospholipase A_2_ (iPLA_2_) VIA gene *CG6718*, the *Drosophila melanogaster* ortholog of human *PLA2G6/PARK14*, mutations in which cause a suite of NDs collectively called *PLA2G6*-associated neurodegeneration (PLAN). Our mutants display age-related loss of climbing ability, a symptom of neurodegeneration in flies. Although phospholipase activity commonly is presumed to underlie iPLA_2_-VIA function, locomotor decline in our mutants is rescued by a transgene carrying a serine-to-alanine mutation in the catalytic residue, suggesting that important functional aspects are independent of phospholipase activity. Additionally, we find that iPLA_2_-VIA knockdown in either muscle or neurons phenocopies locomotor decline with age, demonstrating its necessity in both neuronal and non-neuronal tissues. Furthermore, RNA in situ hybridization shows high endogenous *iPLA_2_-VIA* mRNA expression in adult germ cells, and transgenic HA-tagged iPLA_2_-VIA colocalizes with mitochondria there. Mutant males are fertile with normal spermatogenesis, while fertility is reduced in mutant females. Mutant female germ cells display age-related mitochondrial aggregation, loss of mitochondrial potential, and elevated cell death. These results suggest that iPLA_2_-VIA is important for germline mitochondrial integrity in *Drosophila,* which may be relevant for understanding how PLAN develops.

## Introduction

As global population demographics have shifted toward older age, neurodegenerative disease (ND) has become an increasing health burden worldwide (1). Treatment has been confounded by the fact that loss of neurons in ND, which leads to dementia and reduced motor control, is associated with numerous cytopathologies, including DNA damage and epigenetic changes, mitochondrial and lysosomal dysfunction, Ca^+2^ dysregulation, disrupted RNA and protein homeostasis, as well as inflammation (1, 2). Thus, there is a pressing need to better understand the underlying drivers of ND.

Parkinson’s disease is the second most common ND, affecting ∼1% of individuals over the age of 60 (3). Most cases of Parkinson’s disease are sporadic, presumably arising from a complex interplay between genotype and environment. In some cases, though, environmental toxins have been identified as direct causative agents. For example, contamination of the synthetic opioid MPPP (1-methyl-4-phenyl-4-propionoxy-piperidine) with MPTP (1-methyl-4-phenyl-1,2,3,6-tetrahydropyridine), which is metabolized to MPP^+^ (1-methyl 4-pyridinium) in vivo, caused a cluster of acute cases of parkinsonism amongst opiate users (4). The herbicide paraquat and the pesticide rotenone also have been linked to the disease (5, 6). These toxins inhibit the electron transport chain and promote oxidative stress, suggesting that mitochondrial dysfunction is central, and potentially causative in some cases, to the disease pathology (7, 8).

Several inherited disorders that bear striking resemblance to Parkinson’s disease have permitted identification of genetic contributors, the so-called *PARK* genes (9). Many of these genes encode products that can localize to mitochondria and affect mitochondrial activity and quality control, again suggesting a key role for mitochondrial integrity in the disease and positioning it as a potential therapeutic target (7, 10). *PLA2G6/PARK14* encodes the group 6A calcium-independent phospholipase A_2_ (iPLA_2_-VIA, also called iPLA_2_-β). In 2006, this gene was linked to a group of severe NDs, including infant neuroaxonal dystrophy and neurodegeneration with brain iron accumulation, and three years later was found to be responsible for an autosomal recessive dystonia-parkinsonism (11–14).

The iPLA_2_-VIA enzyme, like other PLA_2_s, hydrolyzes fatty acyl chains from the *sn-2* position of glycerophospholipids (15). Acting within the Lands Cycle of deacylation and reacylation, PLA_2_s promote phospholipid remodeling and repair (16). Acyl chain remodeling seems to be especially important for the mitochondrial phospholipid cardiolipin (CL), which is highly susceptible to oxidative damage due to its proximity to the electron transport chain. In the absence of remodeling, accumulation of damaged CL acyl chains is thought to reduce respiratory function and promote oxidative stress and apoptosis (17, 18). iPLA_2_-VIA can localize to mitochondria in several mammalian cell types and has been implicated in CL remodeling (19–25). Moreover, *PLA2G6* mutant animal models display features of ND along with mitochondrial abnormalities, suggesting the possibility that *PLA2G6-*associated neurodegeneration (PLAN) arises from mitochondrial dysfunction in the absence of CL repair (26–29). However, dramatic CL molecular changes have not been found consistently in *PLA2G6* mutants (26, 27, 30), raising questions of how important iPLA_2_-VIA is for CL remodeling and whether this molecular process underlies PLAN.

iPLA_2_-VIA hydrolyzes other phospholipids in addition to CL and can generate signaling mediators (31). iPLA_2_-VIA also bears near its N-terminus 8-9 ankyrin repeats, which can serve as protein-protein interaction domains, although critical interactors have yet to be characterized (32). It has been observed in a variety of subcellular locations and has been implicated in numerous cellular activities, including cell cycle progression, vesicle trafficking, Ca^+2^ homeostasis, ER stress, and apoptosis (15, 21, 31, 33–35). The relationships between iPLA_2_-VIA’s molecular, cellular, and physiological functions currently are unclear. We investigated *iPLA_2_-VIA* in *Drosophila melanogaster* by determining its endogenous expression pattern and generating a new null mutant. *iPLA_2_-VIA* is expressed ubiquitously at low levels in imaginal tissues. Null mutants are viable and show no synthetic lethality or sterility with two key phosphatidylcholine (PC) metabolizing enzymes. Consistent with a number of recent reports, null *iPLA_2_-VIA* mutants show a striking decline in locomotor ability with age (26, 36, 37), which surprisingly can be rescued by a catalytic-dead iPLA_2_-VIA transgene. Whereas prior reports have focused on neurons, for the purpose of modeling ND, we show here that the locomotor decline can be phenocopied with either neuronal-specific or muscle-specific knockdown in control flies, indicating the importance of this gene in multiple adult tissue types. Furthermore, high endogenous expression is observed in both male and female adult germ cells, and transgenic HA-tagged iPLA_2_-VIA-PB localizes prominently to mitochondria there. Although the mammalian homolog has been implicated in male fertility (38), we observe normal fertility and spermatogenesis in male *iPLA_2_-VIA* mutant *Drosophila*. Still, female mutants have reduced fertility, with age-dependent abnormal mitochondrial aggregation, reduced mitochondrial potential, and elevated cell death in the germline. Taken together, our results demonstrate that iPLA_2_-VIA activity in both neurons and muscles protects from age-related locomotor decline but implicate putative non-catalytic mechanisms in its protective function. We also show that iPLA_2_-VIA plays an important role at mitochondria in the female germline.

## Results

### A null iPLA_2_-VIA mutation does not interact genetically with phospholipid metabolism genes

The *Drosophila melanogaster iPLA_2_-VIA (CG6718)* locus encodes four predicted transcripts and two protein isoforms, which differ by 13 amino acids at their N termini (Fig. 1A). Both *Drosophila* isoforms are highly similar to mammalian iPLA_2_-VIA, with a conserved ankyrin repeat region, catalytic residues, and a 1-9-14 calmodulin binding motif (Fig. 1B, (32), the *Drosophila* PA isoform shares 48% similarity and the PB isoform 47.5% similarity with the shorter human iPLA_2_-VIA protein, and they are 50.4% and 49.9% similar to the longer human isoform, respectively). To generate a genomic mutation in *Drosophila iPLA_2_-VIA,* we excised the P element EY5103 and isolated a 1.4 kb deletion, called *iPLA_2_-VIA^Δ23^*, that spans the transcription initiation site as well as both predicted translation start codons (Fig. 1A). RT-PCR confirmed the absence of full-length *iPLA_2_-VIA* mRNA in our mutant, while the EY5103 insertion line retains a low level of *iPLA_2_-VIA* transcript (Fig. 1C, see also (36)). The null mutants are homozygous and hemizygous viable at room temperature (Table 1) but have reduced lifespans (not shown), as reported in previous studies (26, 36, 39).

**Figure 1.**
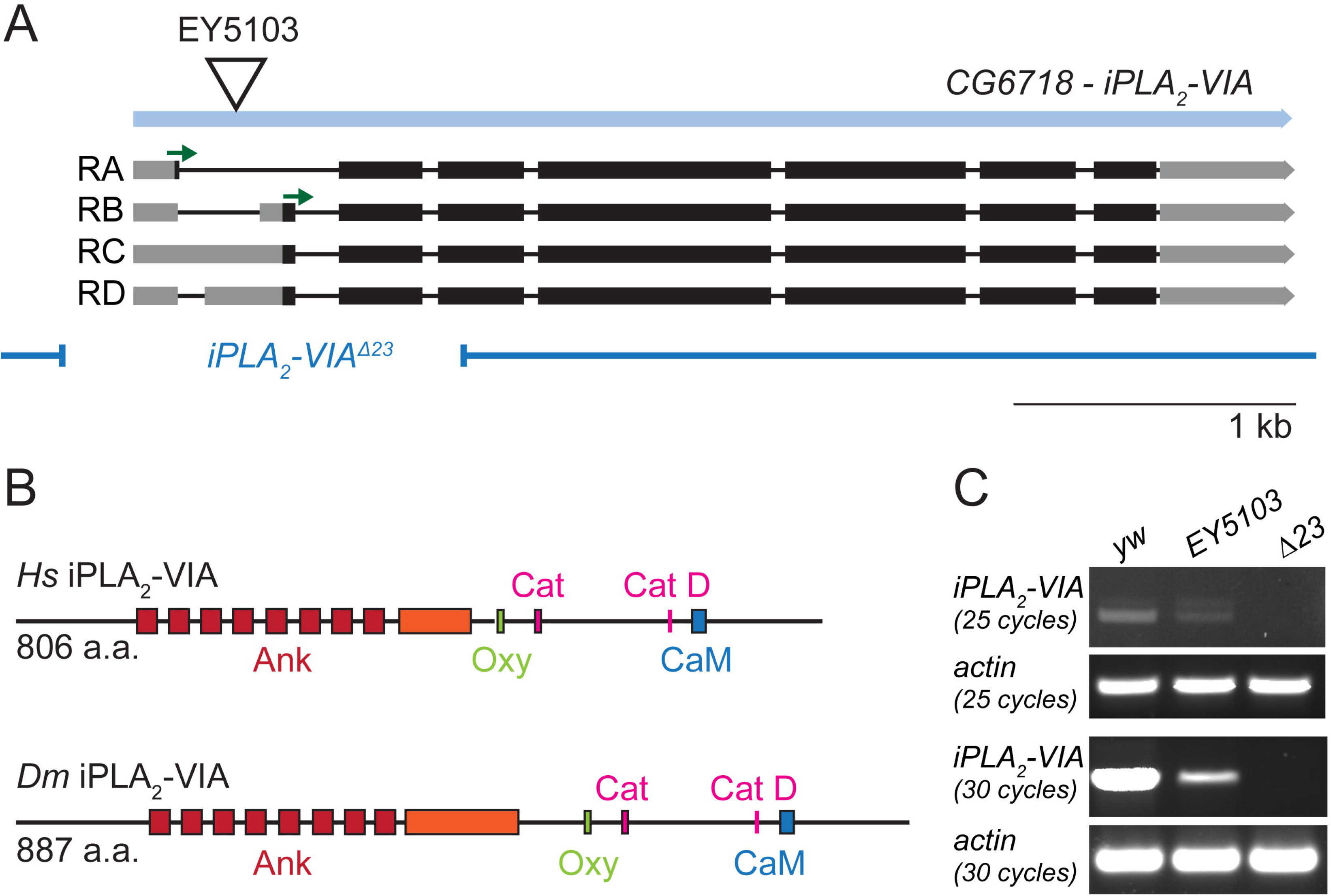
A new mutant allele of *Drosophila iPLA_2_-VIA*. (A) Map of the *CG6718 iPLA_2_-VIA* gene (light blue) showing the insertion site of the EY5103 P element (triangle), predicted transcripts with start codons (green arrows), and the 1.4 kbΔ23 deletion (breakpoints indicated by the dark blue lines). (B) Domain mapping of primary structures demonstrates high homology between *Drosophila* and human iPLA_2_-VIA. Colors are as in (32), with red indicating the ankyrin repeats (Ank; the ninth ankyrin repeat in the shorter mammalian iPLA_2_-VIA isoform is disrupted by additional residues in the fly proteins and in the longer mammalian iPLA2-VIA isoform, indicated here in orange), green indicating the oxyanion hole that coordinates the substrate during catalysis (Oxy), pink indicating the catalytic site (Cat, GTSTG) and catalytic aspartate residue (Cat D), and blue indicating the 1-9-14 calmodulin binding motif (CaM). (C) Whole fly RT-PCR demonstrates the absence of full-length *iPLA_2_-VIA* mRNA in the Δ23 mutant. The EY5103 insertion line retains low levels of *iPLA_2_-VIA* mRNA.

**Table 1.**
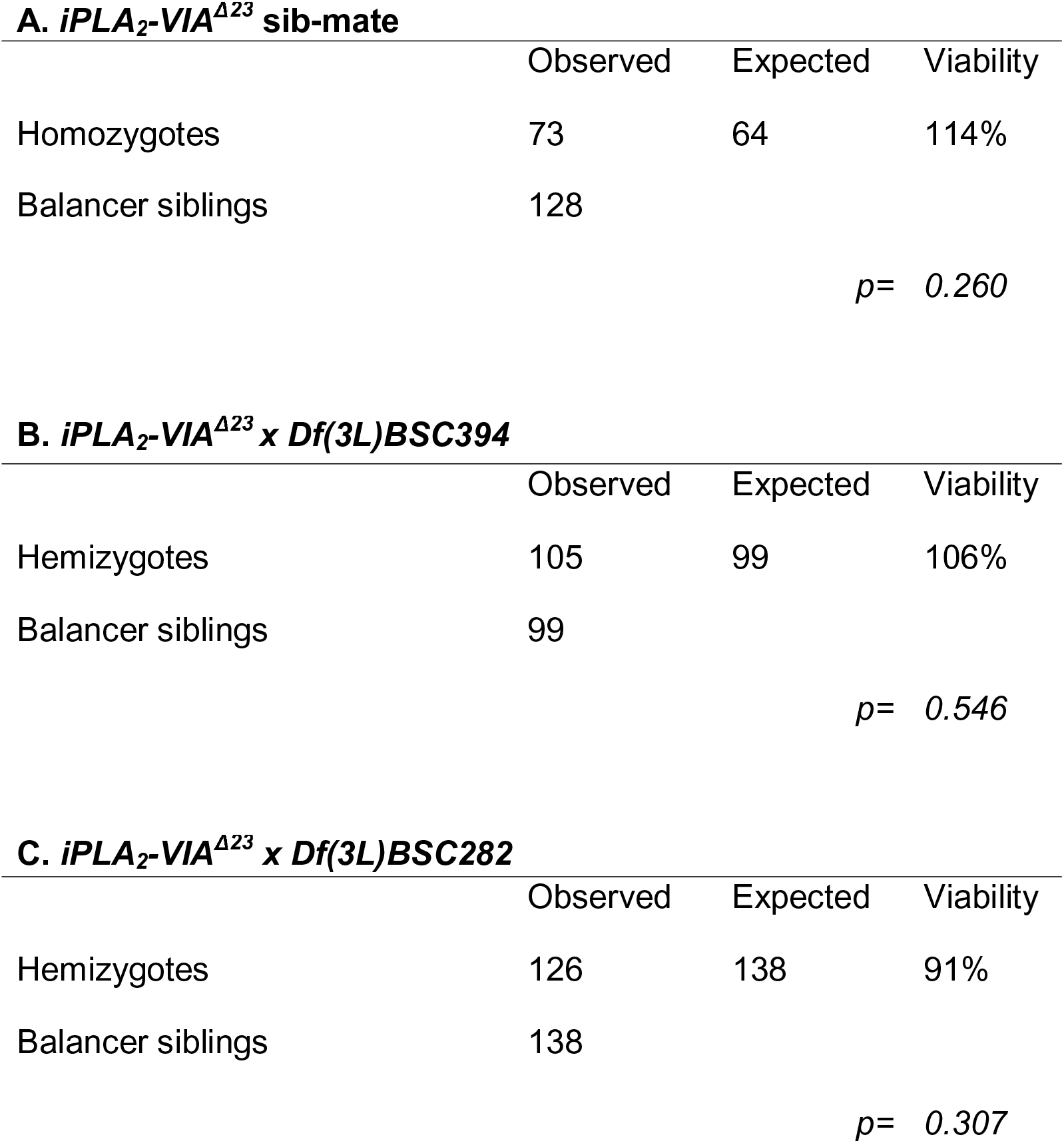
iPLA_2_-VIA^Δ23^ mutants are viable. iPLA_2_-VIA^Δ23^ mutants are homozygous (A) and hemizygous (B, C) viable. Numbers are F1 progeny (“observed”). Expected numbers are derived from the observed number of balancer siblings and the expected Mendelian ratios. “Viability” represents the proportion of observed homozygous or hemizygous F1 flies compared to the expected number. Statistical comparison by chi-square test.

|  | Observed | Expected | Viability |
| --- | --- | --- | --- |
| Homozygotes | 73 | 64 | 114% |
| Balancer siblings | 128 |  |  |
| *p*= 0.260 |  |  |  |

**B. *iPLA<sub>2</sub>-VIA*<sup>Δ23</sup> x *Df(3L)BSC394***
|  | Observed | Expected | Viability |
| --- | --- | --- | --- |
| Hemizygotes | 105 | 99 | 106% |
| Balancer siblings | 99 |  |  |
| *p*= 0.546 |  |  |  |

**Table 1. *iPLA<sub>2</sub>-VIA*<sup>Δ23</sup> mutants are viable.**
|  | Observed | Expected | Viability |
| --- | --- | --- | --- |
| Hemizygotes | 126 | 138 | 91% |
| Balancer siblings | 138 |  |  |
| *p*= 0.307 |  |  |  |

Early studies implicated iPLA_2_-VIA in phospholipid homeostasis (33). For example, overexpressing the phosphatidylcholine (PC) synthesis enzyme CDP phosphocholine cytidylyltransferase (Pcyt1) in mammalian cells led to compensatory upregulation of iPLA_2_-VIA expression and activity, to catabolize the excess PC (40, 41). To explore whether this role is conserved at the organismal level in *Drosophila*, we obtained a Pcyt1 overexpression line, *GS15374,* and examined *iPLA_2_-VIA* mRNA levels by reverse transcription and PCR (RT-PCR). In whole flies, *iPLA_2_-VIA* mRNA expression does not increase when Pcyt1 is overexpressed (Fig. S1A, E). Additionally, no lethality is observed when Pcyt1 is overexpressed in the *iPLA_2_-VIA^Δ23^* mutant (Table 2A), suggesting that iPLA_2_-VIA is not uniquely essential to catabolize excess PC when Pcyt1 is upregulated. We also tested for genetic interactions between our *iPLA_2_-VIA^Δ23^* mutation and *pcyt1* loss of function (42), reasoning that removing *iPLA_2_-VIA* from the *pcyt1* mutant might partially suppress its phenotype by allowing more PC to accumulate. However, neither the lethality nor the sterility of *pcyt1^16919^* mutants is suppressed by the absence of *iPLA_2_-VIA* (Table 2B-D), and iPLA_2_-VIA mRNA levels are not reduced in *pcyt1* mutant flies (Fig. S1B, F). Sws/NTE is a phospholipase in the same family as iPLA_2_-VIA and is known to be important for PC homeostasis (43). Loss of *iPLA_2_-VIA* does not enhance the lethality of *sws^4^* mutants (Table 3), suggesting that iPLA_2_-VIA does not compensate for Sws/NTE in processes that affect viability. Finally, *iPLA_2_-VIA* mRNA expression is unchanged in whole flies mutant for the ethanolamine kinase *easily shocked (eas)*, a key enzyme in phosphatidylethanolamine (PE) synthesis (Fig. S1C, G, (44)). Altogether, our results do not point to a major unique role for *Drosophila* iPLA_2_-VIA in phospholipid housekeeping.

**Table 2.** iPLA_2_-VIA and pcyt1 do not show synthetic effects on viability or fertility. (A) Overexpressing Pcyt1 does not cause lethality in a control background or in the *iPLA_2_-VIA^Δ23^* mutant. Numbers of flies overexpressing Pcyt1 and sibling flies lacking the *tubulin-GAL4* driver are shown. “Viability” represents the proportion of overexpressing flies compared to the matched siblings without the *tubulin-GAL4* driver. Statistical comparison by chi-square test. (B) Although *pcyt1^16919^* homozygotes are sub-viable, *pcyt1^16919^ iPLA_2_-VIA^Δ23^* double mutants show no additional lethality. Numbers shown are F1 progeny (“observed”) from mated heterozygous parents. Expected numbers are derived from the observed number of balancer siblings and the expected Mendelian ratios. “Viability” represents the proportion of observed homozygous F1 flies compared to the expected number. The proportion of *pcyt1^16919^* homozygotes was compared to the proportion of *pcyt1^16919^ iPLA_2_-VIA^Δ23^* double mutant homozygotes using two proportion z-test. (C) Both *pcyt1^16919^* homozygous females and *^pcyt116919^ iPLA2-VIA^Δ23^* doublemutant homozygous females are completely sterile in individual crosses to single *yw* males at 23°C. Crosses were kept and monitored for signs of progeny for 8-10 days. (D) Both *pcyt1^16919^* homozygous males and *pcyt1^16919^ iPLA_2_-VIA^Δ23^* doublemutant homozygous males are markedly sub-fertile in individual crosses to single *yw* females at 23°C. Parental mating pairs were kept for 8-10 days. Average number of progeny from individual crosses shown ± standard deviations, comparison by unpaired t-test.

| **A. *Pcyt1* overexpression** | Observed | Expected | Viability |
| --- | --- | --- | --- |
| *tub-GAL4 > UAS-pcyt1; iPLA<sub>2</sub>-VIA<sup>Δ23</sup>* | 48 | 45 | 107% |
| *UAS-pcyt1; iPLA<sub>2</sub>-VIA<sup>Δ23</sup>* | 45 |  |  |
|  |  |  | *p*= 0.655 |
| *tub-GAL4 > UAS-pcyt1; iPLA<sub>2</sub>-VIA<sup>+</sup>* | 29 | 29 | 100% |
| *UAS-pcyt1; iPLA<sub>2</sub>-VIA<sup>+</sup>* | 29 |  |  |
|  |  |  | *p*= 1.00 |

| **B.** | ***pcyt1<sup>16919</sup> iPLA<sub>2</sub>-VIA<sup>Δ23</sup> sib-mate*** |  |  | ***pcyt1<sup>16919</sup> sib-mate*** |  |  |
| --- | --- | --- | --- | --- | --- | --- |
|  | Observed | Expected | Viability | Observed | Expected | Viability |
| Homozygotes | 29 | 63.5 | 46% | 21 | 41 | 51% |
| Balancer siblings | 127 |  |  | 82 |  |  |
|  |  |  |  |  |  | *p*= 0.720 |

| **C.** | ***pcyt1<sup>16919</sup> iPLA<sub>2</sub>-VIA<sup>Δ23</sup> females***<br>(n = 10) |  | ***pcyt1<sup>16919</sup> females***<br>(n = 9) |
| --- | --- | --- | --- |
| Average number of progeny | 0 |  | 0 |

**Table 2. *iPLA<sub>2</sub>-VIA* and *pcyt1* do not show synthetic effects on viability or fertility.**
| **D.** | ***pcyt1<sup>16919</sup> iPLA<sub>2</sub>-VIA<sup>Δ23</sup> males***<br>(n = 10) |  | ***pcyt1<sup>16919</sup> males***<br>(n = 10) |  |
| --- | --- | --- | --- | --- |
| Average number of progeny | 0.4 ± 0.843 |  | 1.6 ± 2.55 |  |
|  |  |  |  | *p*= 0.185 |

**Table 3.** *iPLA_2_-VIA* and *sws* do not show synthetic loss of viability. Although *sws_4_* mutants are sub-viable, *sws_4_ iPLA_2_-VIA^Δ23^* double mutants show no additional lethality. Numbers shown are F1 progeny (“observed”) from mated heterozygous parents (male parents are heterozygous for *iPLA_2_-VIA^Δ23^* but hemizygous for *sws_4_*). Expected numbers are derived from the observed number of balancer siblings and the expected Mendelian ratios. “Viability” represents the proportion of observed mutant F1 flies compared to the expected number. Proportion of *sws_4_ iPLA_2_-VIA^Δ23^* double mutants versus *iPLA_2_-VIA^Δ23^* single mutants was compared to the proportion of sibling *sws_4_* mutants versus balancer controls using two proportion z-test.

|  | *sws*<sup>4</sup> *iPLA*<sub>2</sub>-*VIA*<sup>Δ23</sup> sib-mate |  |  |
| --- | --- | --- | --- |
|  | Observed | Expected | Viability |
| Double mutant | 26 | 34 | 76% |
| *iPLA*<sub>2</sub>-*VIA* mutant | 34 |  |  |
| *sws* mutant | 70 | 81 | 86% |
| Balancer siblings | 81 |  |  |
|  |  |  | *p*= 0.691 |

### iPLA_2_-VIA in neurons and muscles maintains locomotor ability with age, via a partially catalytic-independent activity

Because *iPLA_2_-VIA* is associated with ND in humans, we tested our mutant flies for their climbing ability, which is known to decline with age under conditions of neurodegeneration (45). In accord with other reports, *iPLA_2_-VIA^Δ23^* mutants exhibit climbing defects after 20 days of age, in contrast to background control and isogenic control animals (Fig. 2A-B, light bars (26, 36, 39)). Ubiquitous expression of a C-terminally HA-tagged iPLA_2_-VIA-PB wild-type cDNA transgene rescues the climbing ability of the mutant, confirming the locus-specificity of the defect (Fig. 2C-D, dark bars). Surprisingly, a transgene in which the catalytic serine is replaced with an alanine residue also rescues the climbing defect of mutant flies, although to a weaker extent than the wild-type transgene, suggesting that iPLA_2_-VIA function is partially independent of catalytic activity (Fig. 2C-D, yellow bars, (32)).

Ubiquitous RNAi knockdown of *iPLA_2_-VIA* using *tubulin-GAL4* phenocopies the mutant in both males (Fig. 2E, gray bars) and females (Fig. 2F, gray bars). Knocking down *iPLA_2_-VIA* only in neurons with the *elav-GAL4* driver produces a climbing defect that is markedly weaker than in the ubiquitous knockdown (Fig. 2G-H), suggesting a possible requirement in additional tissues. In accord with this, RNAi knockdown of *iPLA_2_-VIA* with the muscle-specific driver *DJ667-GAL4* also reproduces the climbing defect (Fig. 2I-J, gray bars). In comparison to knockdown of *pink1*, a gene required for maintenance of muscle integrity with age (Fig. 2I-J, red bars, (46, 47)), *iPLA_2_-VIA* knockdown is comparable at 20 days in males and even stronger by 30 days of age in both males and females. Thus, iPLA_2_-VIA is required in neurons and muscle, and possibly other tissues, for maintenance of normal locomotor activity with age. Publicly available high throughput transcriptomic data show moderate to strong expression in gut, fat body, and heart, all tissues known to affect aging (flybase.org; (48)).

**Figure 2.**
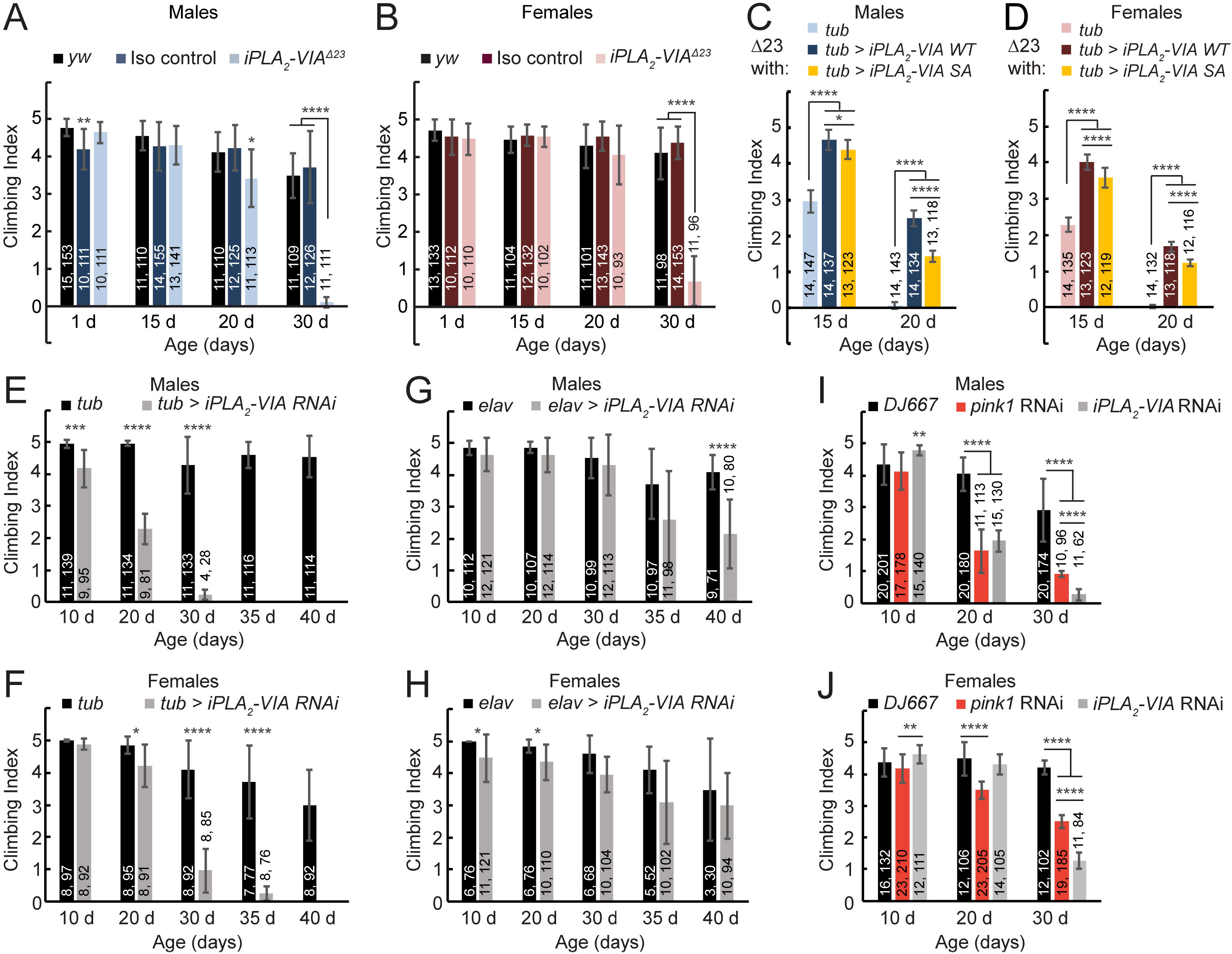
Impaired climbing ability in aged *iPLA_2_-VIA^Δ23^* mutants can be rescued by a catalytic dead *iPLA_2_-VIA-PB* transgene and phenocopied with neuronal or muscle knockdown. (A-B) Male (A, light blue) and female (B, light pink) *iPLA_2_-VIA^Δ23^* mutant adults have normal climbing ability until 20 days of age at room temperature. After 20 days, climbing ability declines compared to *yw* (black) and isogenic controls (dark blue in A, and dark red in B). Newly eclosed isogenic control males (A, dark blue) show slightly reduced climbing ability compared to *yw* males. (C-D) *Tubulin-GAL4* driven expression of wild-type *UAS-iPLA_2_-VIA-HA* in males (C, dark blue bars) and females (D, dark red bars) at 26°C rescues the climbing defect of aged *iPLA_2_-VIA^Δ23^* mutants at 15 and 20 days of age. 26°C was used to increase GAL4 activity in rescue and knockdown experiments. Note that mutant flies show climbing defects by 15 days at this higher temperature. In this experiment, mutants carry the GAL4 driver without the resue transgene (light blue bars in C, and light pink bars in D). *Tubulin-GAL4* driven expression of a *UAS-iPLA_2_-VIA-SA* transgene carrying a serine-to-alanine mutation in the catalytic residue at 26°C also partially rescues the climbing defect (yellow bars), although rescue is weaker than with the wild-type transgene. Wild-type and SA transgenes were transformed into the same genomic locus using the phi-C31 system. (E-F) Ubiquitous expression of *UAS-*driven double stranded RNA targeting *iPLA_2_-VIA* (*HMS01544*) with *tubulin-GAL4* at 26°C phenocopies the climbing defect in male (E) and female (F) flies (gray bars, compared to *GAL4-*only controls, black bars). Knockdown flies die by 35 and 40 days of age in males and females, respectively. (G-H) Expressing *HMS01544* in neurons only with *elav-GAL4* at 26°C weakly phenocopies the climbing defect in male (G) and female (H) flies (gray bars, compared to *GAL4-*only controls, black bars). (I-J) Expressing *HMS01544* in muscles only with *DJ667-GAL4* produces a strong phenocopy of the age-induced climbing defect in male (I) and female (J) flies, similar to that resulting from knockdown of *pink1 (HMS02204,* red bars). Note that *pink1* knockdown flies also have moderately reduced climbing ability at 10 days of age compared to *iPLA2-VIA* knockdown flies, but this was not statistically significant compared to GAL4-only controls. Number of groups (first number) and number of flies (second number) assayed for each condition is shown on the graphs. Bars represent the average climbing index for each condition. Error bars are standard deviations. Differences assessed by unpaired t-test, *p < 0.05, **p < 0.01, ***p < 0.005, ****p < 0.0005.

### Drosophila iPLA_2_-VIA is expressed strongly in adult germline but is not required for spermatogenesis

To explore further the endogenous expression of *Drosophila iPLA_2_-VIA,* we performed RNA in situ hybridization. We observe weak mRNA expression in wild-type imaginal tissues and no detectable expression in wild-type larval brains (Fig. 3A-F). Strikingly, strong expression is seen in the germ cells of both male and female adults Fig. 3G-H). Because iPLA_2_-VIA has been implicated previously in mammalian male fertility (38), we examined male fertility in our mutants. Surprisingly, *iPLA_2_-VIA^Δ23^* mutant males are fertile, even when they are aged (Fig. 4A-B).

**Figure 3.**
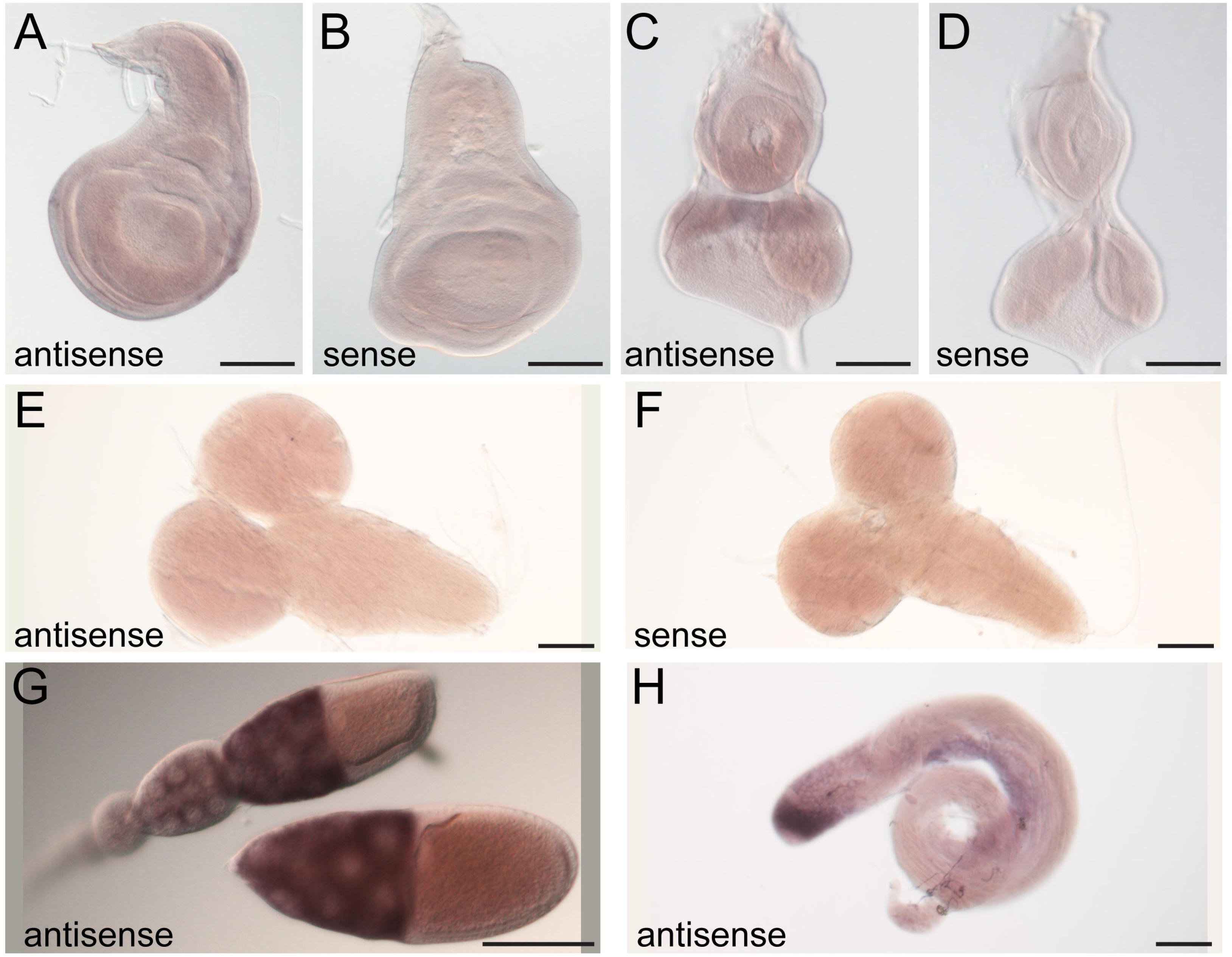
*iPLA_2_-VIA* mRNA is highly expressed in the adult germline. In situ hybridization to endogenous *iPLA_2_-VIA* mRNA (purple) in *w^1118^* animals shows weak ubiquitous expression in wild-type wing imaginal discs (A) and eye-antennal discs (C). Sense probes (B, D) were used as controls. Expression is negligible in larval brains (E, compare to sense probe control in F). Strong expression is seen in both female (G) and male (H) adult germlines. Scale bars: 100 μm.

Loss of function mutations in the enzyme Tafazzin (Taz), a mitochondrially-localized transacylase responsible for most of the CL remodeling activity in cells, lead to male infertility with defects in spermatid individualization in *Drosophila* (23, 49). Previous results implicated iPLA_2_-VIA in this pathway, with reduced *iPLA_2_-VIA* expression compensating for loss of *taz* in male fertility (23). We therefore closely examined several aspects of spermatogenesis in *taz*^Δ^*^161^* or *iPLA_2_-VIA^Δ23^* single mutants, as well as in *taz^Δ161^; iPLA_2_-VIA^Δ23^* double mutants. In both single mutants and the double mutant, developmentally programmed mitochondrial aggregation is normal in post-meiotic spermatids (Fig. 4C-F), and mitochondrial derivatives begin elongation normally (Fig. 4G-J). At the individualization stage, the actin-rich individualization complexes (ICs) can be detected by phalloidin staining (Fig. 4K, (50)). No IC defects are seen in *iPLA_2_-VIA^Δ23^* mutants (Fig. 4L). In contrast, ICs are highly disorganized in *taz*^Δ^*^161^* single mutants (Fig. 4M, (23)) and in *taz*^Δ^*^161^; iPLA_2_-VIA^Δ23^* double mutants (Fig. 4N), with no rescue of the *taz* mutant phenotype by removal of *iPLA_2_-VIA* (Fig. 4O). We also tested the fertility of both *taz*^Δ^*^161^*; *iPLA_2_-VIA^Δ23^* double mutant males (n = 8) and *taz*^Δ^*^161^*; *iPLA_2_-VIA^EY5103^* double mutant males (n = 5) in individual crosses to *yw* females and found both to be completely sterile (0 progeny produced over the course of five days). Therefore, as the *iPLA_2_-VIA* mutant neither shows similar defects to the *taz* mutant nor rescues it, our data do not support the idea that iPLA_2_-VIA is a critical unique player in CL remodeling during *Drosophila* spermatogenesis.

**Figure 4.**
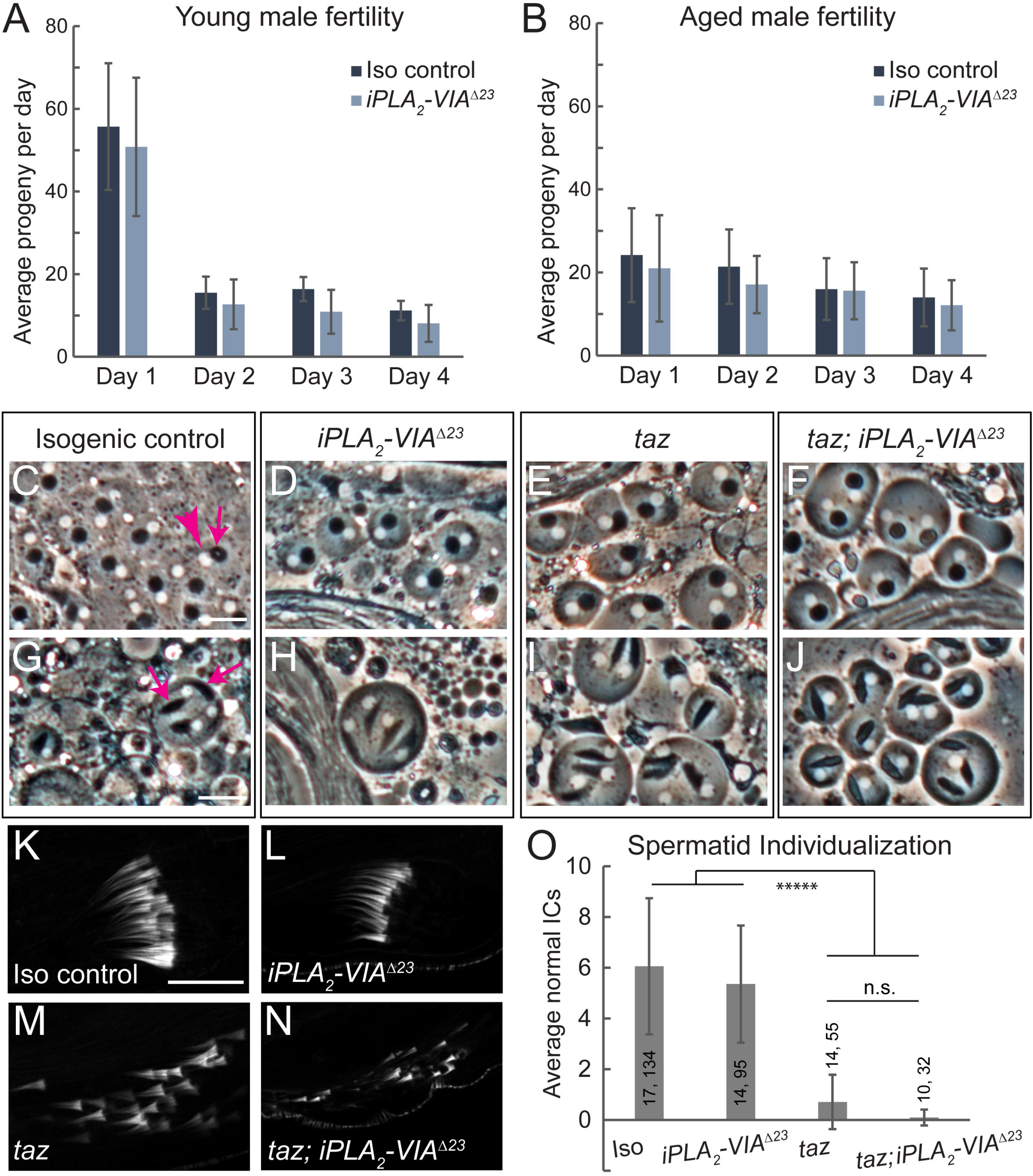
Male fertility and spermatogenesis are normal in *iPLA_2_-VIA^Δ23^* mutants. (A-B) Young (A, 4-5 day old) and aged (B, 20-21 day old) *iPLA_2_-VIA^Δ23^* mutant males (light blue bars) show no fertility defect compared to isogenic controls (dark blue bars). Bars depict average number of adult progeny produced by 10 individual males over the course of four days, error bars represent standard deviation. (C-J) Testis squashes reveal normal post-meiotic mitochondrial Nebenkerne (phase-dark structures, indicated by the magenta arrow in C) in *iPLA_2_-VIA^Δ23^* mutants (D), *taz*^Δ*161*^ mutants (E), and *taz*^Δ*161*^ *iPLA_2_-VIA^Δ23^* double mutants (F), and each Nebenkern is paired with a similarly sized nucleus (phase-light structures, indicated by the magenta arrowhead in C). In slightly later spermatogenic cysts (G-J), mitochondrial derivatives (indicated by magenta arrows in G) elongate normally in all genotypes. Scale bars: 20 μm. (K-N) Phalloidin staining reveals the individualization complexes (ICs) of the maturing spermatids. In *iPLA_2_-VIA^Δ23^* mutants, ICs are normal (L), in contrast with *taz*^Δ*161*^ mutants, in which ICs are highly disorganized (M). *taz*^Δ*161*^ *iPLA_2_-VIA^Δ23^* double mutants show disorganized ICs (N), like *taz*^Δ*161*^ single mutants. Scale bar: 20 μm. (O) IC quantification shows no rescue of the *taz*^Δ*161*^ individualization phenotype by *iPLA_2_-VIA^Δ23^* mutation. Number of testes (first number) and number of individualization complexes (second number) quantified for each genotype are indicated on the graph. Bars depict the average number of normal ICs for each genotype, error bars represent standard deviation. Differences assessed by unpaired t-test, *****p < 10^-5^, n.s. indicates not significant.

### Drosophila iPLA_2_-VIA mutants have reduced female fertility with abnormal mitochondrial distribution in the germline

Consistent with strong *iPLA_2_-VIA* expression in the female germline (Fig. 3G), there is a significant decrease in female fertility in *iPLA_2_-VIA^Δ23^* homozygotes (Fig. 5A), and the same effect is seen in hemizygotes, supporting the locus-specificity of the phenotype (Fig. 5B). This is accompanied by a decrease in number of eggs laid (Fig. 5C-D) but not maternal lethality in embryos or larvae (Fig. S2), suggesting that *iPLA_2_-VIA* is necessary before fertilization, during oogenesis. Although major developmental defects are not evident in *iPLA_2_-VIA^Δ23^* mutant ovaries, we observe striking mitochondrial aggregation in the germlines of aged mutant females. The mitochondrially targeted fluorescent reporter transgene *Psqh-mito-EYFP*, which consists of YFP fused to the mitochondrial localization signal of human cytochrome c oxidase 8A, decorates a diffuse mitochondrial network in the nurse cells of young females, in both null mutants and controls (Fig. 6A-B). By three weeks of age, mitochondria aggregate abnormally in the mutant, despite maintaining their normal diffuse localization in background control and heterozygous genotypes (Fig. 6C-D). Aggregation is especially noticeable in mid-oogenesis stages 8-9. In younger egg chambers, mitochondria are less diffuse even in control genotypes, making the mutant phenotype harder to distinguish (for example, see Fig. 7A’). To quantify mitochondrial aggregation, we used ImageJ to outline the mito-YFP signal areas. In egg chambers with a normal diffuse mitochondrial network, the outlined mito-YFP signal is extensive and complex, but in egg chambers with aggregated mitochondria, the outlines separating areas of signal from background are sparser (Fig. 6E-F). Taking the raw integrated density of the mito-YFP outlines in the area of the nurse cells in ImageJ reveals a highly significant quantitative reduction in the intricacy of the mito-YFP network in mutant germlines at mid-oogenesis compared to controls, in females three weeks of age and older (Fig. 6G-H). Although *iPLA_2_-VIA^Δ23^* mutant flies do not survive to six weeks of age (26, 36, 39), neither control genotype nor mito-YFP stock homozygotes show mitochondrial aggregation even at this late age (Fig. 6J, Fig. S3A-B).

**Figure 5.**
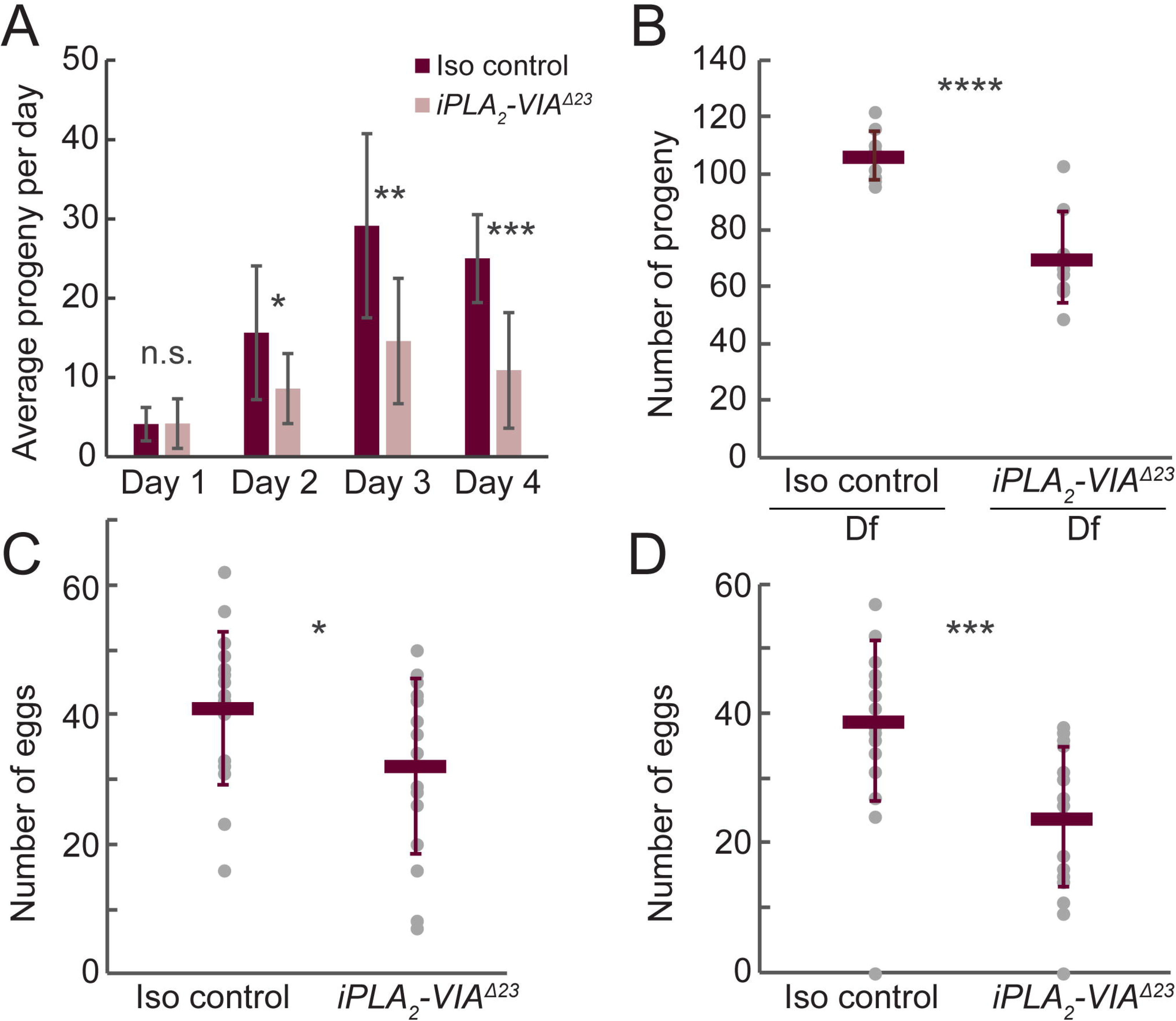
Female fertility is reduced in *iPLA2-VIA^Δ23^* mutants. (A) Young (<1 week old) *iPLA_2_-VIA^Δ23^* homozygous mutant females (light pink bars) produce fewer adult progeny compared to isogenic controls (dark red bars). Bars depict average number of adult progeny produced by 10 individual females over the course of four days, error bars represent standard deviations. (B) Hemizygous mutant females (*iPLA_2_-VIA^Δ23^/Df(3L)BSC282)* produce fewer adult progeny compared to isogenic controls. Dark red bars represent average number of adult progeny produced by 10 individual females over the course of four days, error bars are standard deviations. (C-D) Egg laying by *iPLA_2_-VIA^Δ23^* mutants is reduced compared to isogenic controls in young (C, one week old) and aged (D, 18-19 day old) females. Dark red bars represent average number of eggs produced by 20 individual females over the course of four days, error bars are standard deviations. Differences assessed by unpaired t-test, n.s. not significant, *p < 0.1, **p < 0.02, ***p < 0.001, ****p < 0.0001.

**Figure 6.**
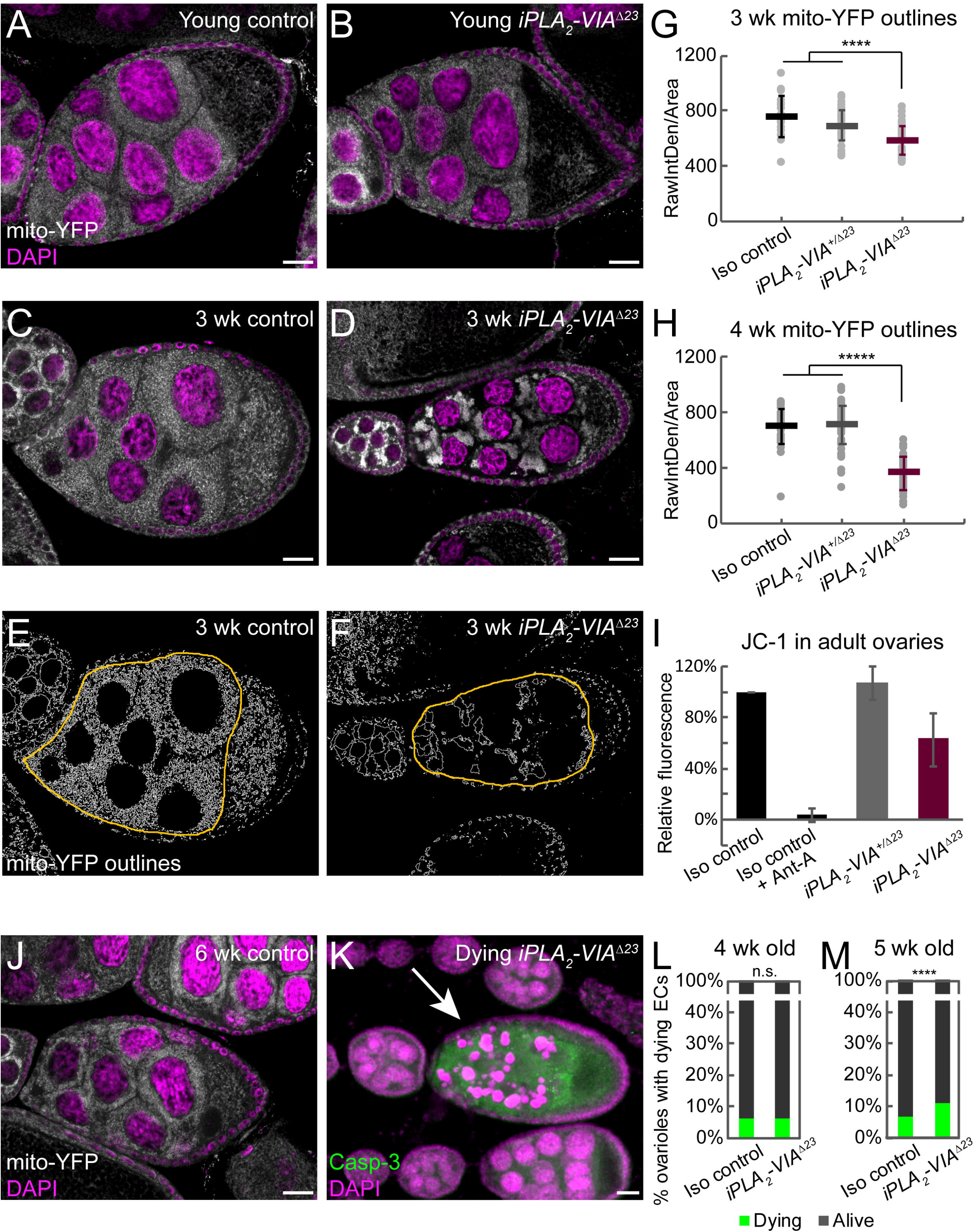
Germ cells from *iPLA2-VIA^Δ23^* mutant females have abnormal mitochondrial distribution, reduced mitochondrial potential, and elevated apoptosis. (A-B) *Psqh-mito-EYFP* (white) was used to observe the mitochondrial distribution in female germ cells from young (<1 week old) *iPLA_2_-VIA^Δ23^* mutants (*Psqh-mito-EYFP, iPLA_2_-VIA^Δ23^/iPLA_2_-VIA^Δ23^)* and heterozygous controls (*Psqh-mito-EYFP, iPLA_2_-VIA^Δ23^/revertant*Δ*11*). Nuclei are stained with DAPI (magenta). (C-D) Mito-YFP labeled mitochondria appear clumpy in germ cells from aged (3 week old) *iPLA_2_-VIA^Δ23^* female flies (D) but not controls (C, *Psqh-mito-EYFP/revertant*Δ*11*; for *Psqh-mito-EYFP, iPLA_2_-VIA^Δ23^/revertant*Δ*11* heterozygotes see quantification in G and H). (E-H) Mito-YFP signal within the area of the germline nurse cells was outlined using ImageJ (E-F, yellow outlines) and the raw integrated density of the outline signal was used to quantify clumpiness (G-H). Homozygous *iPLA_2_-VIA^Δ23^* mutants (dark red, *Psqh-mito-EYFP, iPLA_2_-VIA^Δ23^/iPLA_2_-VIA^Δ23^*, n=39) show significantly reduced mito-YFP outline signal compared to heterozygotes (gray, *Psqh-mito-EYFP, iPLA_2_-VIA^Δ23^/revertant*Δ*11,* n=43) and isogenic controls (black, *Psqh-mito-EYFP/revertant*Δ*11,* n=19) at 3 weeks (G) and 4 weeks old (H; black, *Psqh-mito-EYFP/revertant*Δ*11,* n=42; gray, *Psqh-mito-EYFP, iPLA_2_-VIA^Δ23^/revertant*Δ*11,* n=67; dark red, *Psqh-mito-EYFP, iPLA_2_-VIA^Δ23^/iPLA2-VIA^Δ23^*, n=50), indicating increased clumpiness. Bars represent averages and standard deviations for each condition. Differences assessed by unpaired t-test, ****p < 0.0001, *****p < 10^-6^. (I) Ovaries from 4 week old *iPLA_2_-VIA^Δ23^* females (dark red bar) show reduced JC-1 fluorescence compared to age matched controls (black, *Psqh-mito-EYFP/revertant*Δ*11;* gray, *Psqh-mito-EYFP, iPLA_2_-VIA^Δ23^/revertant*Δ*11*) using plate-based fluorimetry. In each experiment, the red (595 nm) JC-1 fluorescence of the heterozygote or the mutant is expressed a s a percentage of the fluorescence of the control, and 3-4 biological replicates are averaged. Error bars are standard deviations. Antimycin A was used as a mitochondrial poison to demonstrate the specificity of the JC-1 signal. (J) Controls (*Psqh-mito-EYFP/revertant*Δ*11*) do not show mito-YFP clumpiness even at 6 weeks of age, while *iPLA_2_-VIA^Δ23^* mutant females do not survive to 6 weeks. (K) By 5 weeks of age, *iPLA_2_-VIA^Δ23^* mutants show increased levels of germline apoptosis, marked by cleaved caspase-3 staining (green) and fragmentation of the nurse cell nuclei (DAPI, magenta), quantified in M. (L-M) The percentage of ovarioles containing dying egg chambers (green bars) is elevated in *iPLA_2_-VIA^Δ23^* mutants compared to isogenic controls at 5 weeks of age (M, >1300 ovarioles counted for each genotype across four biological replicates) but not 4 weeks (L, >900 ovarioles counted for each genotype across three biological replicates). Dying egg chambers were identified by their fragmented nurse cell nuclei and cleaved caspase-3 staining, as shown in (K). Statistical comparison by two proportion z-test, ****p < 0.0001. Scale bars: 20 μm.

**Figure 7.**
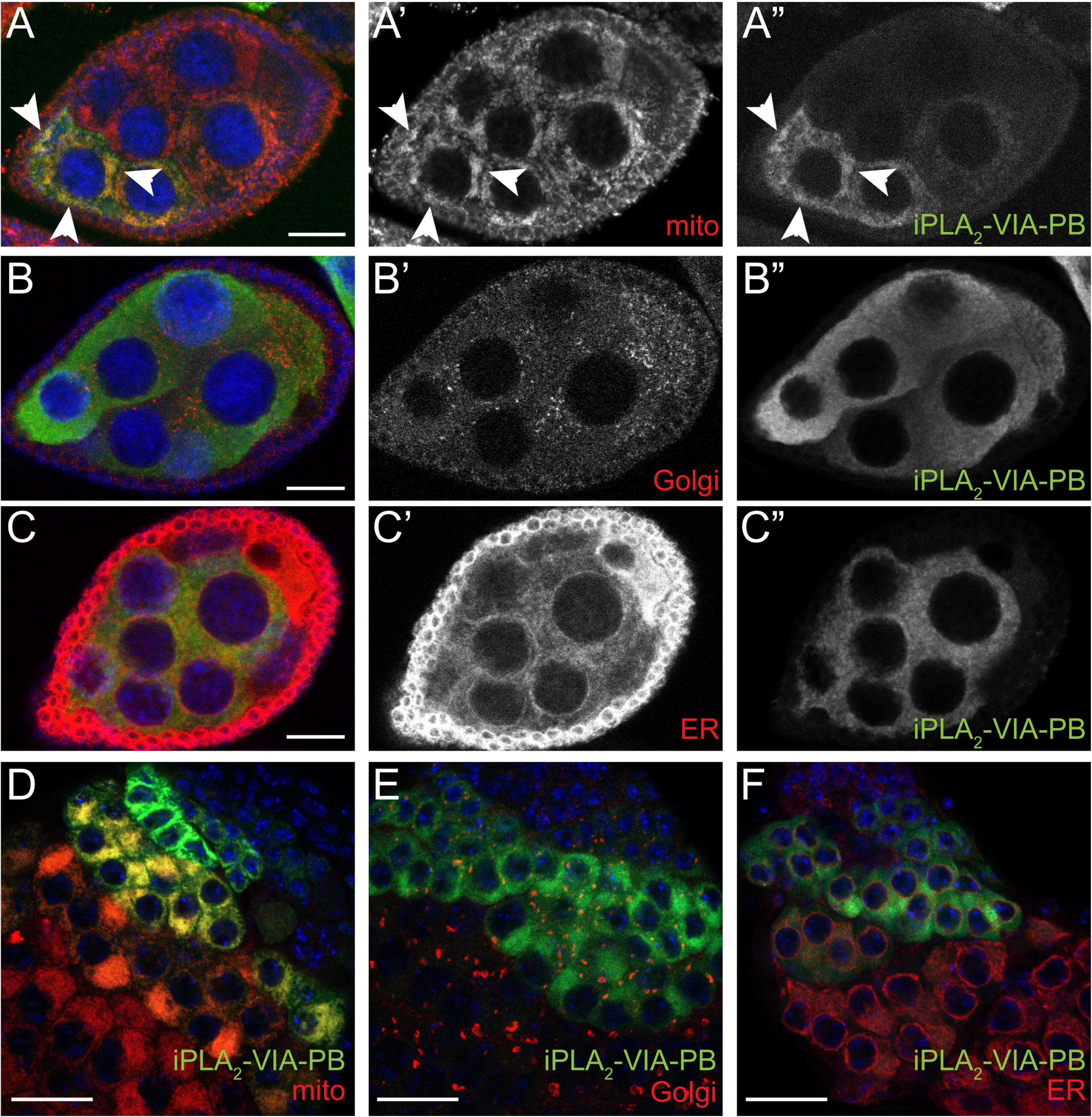
iPLA2-VIA-PB protein localizes to mitochondria in the female and male germ cells. (A-C) In the female germline, HA-tagged wild-type iPLA_2_-VIA-PB (green, grayscale shown in A”-C”, expressed with *NGT40-GAL4*) strongly colocalizes with a mitochondrial marker (A, red, *Psqh-mito-EYFP*, arrowheads) and shows minimal colocalization with Golgi (B, red, anti-Golgin 84) and ER (C, red, anti-Calnexin 99A) markers. Individual channels for mitochondria, Golgi, and ER shown in A’, B’, C’, respectively. (D-F) In male germ cells, iPLA_2_VIA-PB (green, expressed with *bam-GAL4-VP16*) also colocalizes more strongly with a mitochondrial marker (D, red, *UAS*746 *mCherry-mitoOMM*) than Golgi (E, red) or ER (F, red) markers. Scale bars: 20 μm.

Because mitochondrial aggregation can result from damage (51, 52), we examined mitochondrial potential in dissected ovaries using the potential-sensitive fluorescent dye JC-1. *iPLA_2_-VIA^Δ23^* mutant ovaries have reduced JC-1 fluorescence compared to controls, indicating loss of mitochondrial potential (Fig. 6I). By five weeks of age, germ cells of the mutant undergo apoptosis, with fragmented nuclei and cleaved caspase-3 staining, in contrast to controls (Fig. 6K-M).

To confirm further that mitochondrial aggregation is independent of the mito-YFP transgene, we examined another mitochondrial marker, immunofluorescence for the ATP-5A protein. We again saw mitochondrial aggregation in germ cells from aged *iPLA_2_-VIA ^23^* mutant females and not in those from isogenic control flies (Fig. S3C-D), but we were not able to quantify it due to the variability of the staining. We noticed, though, that on average germ cells from aged *iPLA_2_-VIA ^23^* mutant females have lower ATP-5A signal intensity than those from control flies (Fig. S3E-F), which might indicate that damaged mitochondria in the mutant are subject to quality control mechanisms involving degradation. We do not know why the mito-YFP marker appears more stable than the endogenous ATP-5A protein.

### iPLA_2_-VIA-PB localizes to mitochondria in germ cells

Because iPLA_2_-VIA has been implicated in many different activities in a variety of cellular compartments, we generated an HA-tagged iPLA_2_-VIA wild-type cDNA transgene, PB isoform. After confirming that the transgene is active (Fig. 2C-D), we examined its subcellular localization. In female germ cells, the transgene colocalizes most strongly with a mitochondrial marker (Fig. 7A-C), consistent with the mitochondrial defect in the mutant. The transgene also colocalizes strongly with a mitochondrial marker in male germ cells (Fig. 7D-F). We additionally examined localization in somatic cells using the large polyploid cells of the larval fat body and salivary gland (53). Interestingly, colocalization in these tissues is strongest with a Golgi marker (Fig. S4A, D), suggesting that subcellular localization of iPLA_2_-VIA is regulated differently in distinct cell types.

## Discussion

### iPLA_2_-VIA in phospholipid metabolism

Tight regulation of iPLA_2_-VIA levels and activity is important for cell cycle dependent phospholipid accumulation (54) and PC homeostasis (40, 41) in several mammalian cultured cell lines. However, genetic perturbation of the rate-limiting PC synthesis enzyme Pcyt1 in whole flies does not cause obvious changes in *iPLA_2_-VIA* mRNA expression (Fig. S1), nor do we find evidence of genetic interactions between *iPLA_2_-VIA* and *pcyt1* in viability or fertility (Table 2). To our knowledge, this is the first report on a possible genetic interaction between *iPLA_2_-VIA* and *pcyt1* in an intact organism. *iPLA_2_-VIA* also shows no genetic redundancy for viability with *sws/NTE,* a conserved paralog important for PC homeostasis (Table 3, (43)). We see no global downregulation of *iPLA_2_-VIA* mRNA in an *eas* mutant with reduced PE levels either (Fig. S1). *iPLA_2_-VIA* mRNA expression is very low in wild-type imaginal discs (Fig. 3), despite substantial cell proliferation there (55), and *iPLA_2_-VIA* null mutants are developmentally viable (Table 1). Altogether, our data suggest that iPLA_2_-VIA does not play unique roles in phospholipid accumulation in *Drosophila*, consistent with recent mass spectrometry analysis of another *iPLA_2_-VIA* knockout mutant (39).

### iPLA_2_-VIA in male fertility and CL remodeling

We expected that, like the mouse ortholog, *Drosophila* iPLA_2_-VIA would be important for male fertility, in accord with its high expression in the male germline (Fig. 3, (38)). Furthermore, it was reported to interact genetically with the mitochondrially-localized CL remodeling enzyme Taz in *Drosophila* male fertility (23). We observed iPLA_2_-VIA colocalization with mitochondria in male germ cells (Fig. 7), as has been documented in cultured mammalian cells (19, 21). Surprisingly, however, *iPLA_2_-VIA* mutant males are fully fertile, with normal spermatid mitochondrial morphogenesis and, unlike the *taz* mutant, normal spermatid individualization (Fig. 4). Moreover, we find no evidence that *iPLA_2_-VIA* loss of function mutations interact genetically with *taz,* in contrast to a prior report (23). Our genetic data in the testis therefore may argue against the idea that iPLA_2_-VIA is a major unique player in CL remodeling. This is consistent with mass spectrometry results showing minimal CL changes in *iPLA_2_-VIA* hypomorphic mutant heads or whole flies (26, 30) and with recent studies indicating that Taz can remodel CL independently of phospholipases (56).

### iPLA_2_-VIA in female fertility

*iPLA_2_-VIA* mRNA is highly expressed in the wild-type female germline (Fig. 3G), and *iPLA_2_-VIA* mutant females have reduced fertility compared to controls (Fig. 5). Despite normal ovariole morphology, germ cells in aged *iPLA_2_-VIA* mutant females have abnormally aggregated mitochondria with decreased membrane potential, and they eventually die by apoptosis (Fig. 6). In HeLa cells, pharmacological disruption of mitochondrial membrane potential induces aggregation via stabilization of the mitochondrially targeted kinase PINK1 and recruitment of its substrate and partner Parkin, an E3 ubiquitin ligase. Ubiquitinated Parkin substrates on the outer mitochondrial surface recruit the autophagy receptor p62/SQSTM1, which promotes aggregation, possibly for sequestration before degradation by mitophagy (51, 52). Intriguingly, mitochondrial aggregation in *Drosophila* female germ cells has been observed in *pink1* and *parkin* mutants, similarly to *iPLA_2_-VIA* mutants (57, 58). This might suggest a parallel mechanism for mitochondrial damage detection and aggregation, independent of the PINK1-Parkin pathway, that perhaps relies on another mitophagy receptor, for example BNIP3, which is known to be active in the female germline (59, 60). It is tempting to propose that in the absence of iPLA_2_-VIA, PINK1, or Parkin, mitochondrial damage accumulates and leads to aggregation. Eventually, persistent mitochondrial damage may overwhelm cytoprotective mechanisms like aggregation and mitophagy, leading to apoptosis. We thus hypothesize that iPLA_2_-VIA protects germline mitochondria from accumulated injury that can result in apoptosis, as in other contexts (19, 22, 25). A direct effect is supported by the localization of iPLA_2_-VIA to mitochondria in germ cells (Fig. 7). Reduced mitochondrial membrane potential also has been observed in brains of an *iPLA_2_-VIA* hypomorph (26), possibly suggesting a mechanistic connection between the germline and neuronal phenotypes

### iPLA_2_-VIA in neuromuscular degeneration

Our *iPLA_2_-VIA* mutants show a severe decline in locomotor ability with age, like other *Drosophila* and mouse mutant models (Fig. 2, (26, 36, 37, 61–63)). Phenocopy by RNAi knockdown in neurons is consistent with the idea that iPLA_2_-VIA acts autonomously to protect against neurodegeneration (26, 36). Our data suggest that iPLA_2_-VIA also is important in other tissues to prevent locomotor decline, notably muscle, in accord with a report of muscle degeneration in two human PLAN patients (64). The strength of the phenocopy with muscle-specific knockdown is comparable to that seen for *pink1* (Fig. 2)*. PINK1* and its partner *parkin* are well-established disease loci for autosomal recessive parkinsonism in humans (10). As discussed above, PINK1 accumulates on the outer mitochondrial membrane under stress conditions and recruits Parkin to activate quality control mechanisms, including mitophagy, that maintain the overall integrity of the cellular mitochondrial network (65). In their absence, accumulation of damaged mitochondria leads to degeneration of neurons, as well as muscle in *Drosophila* (46, 47). The parkinsonism in human patients, as well as the locomotor defects and germline mitochondrial aggregation in flies, suggest some similarities between loss of *iPLA_2_-VIA* and loss of *pink1* or *parkin* (57, 58). Notably, however, *Drosophila pink1* and *parkin* mutants display locomotor defects at earlier time points, as well as other indications of muscle degeneration, including crushed thoraces and wing posture abnormalities, not seen in *iPLA_2_-VIA* mutants (46, 47). Additionally, the normal fertility and spermatid mitochondrial morphology in *iPLA_2_-VIA* mutant males stand in contrast to the highly penetrant male germline defects of *pink1* and *parkin* mutants (46, 47, 66).

Although many studies have implicated iPLA_2_-VIA in mitochondrial maintenance via catalytic removal of damaged phospholipid acyl chains, our data show that a transgene with a serine-to-alanine mutation in the catalytic site can rescue the climbing defect of the mutant, consistent with another report in which a human iPLA_2_-VIA transgene carrying the analogous catalytic site mutation rescued bang-sensitivity and lifespan defects of *iPLA_2_-VIA* mutants (39). This suggests that phospholipase activity is not the only important aspect in iPLA_2_-VIA mediated cytoprotection, possibly explaining the observation that mutations associated with dystonia-parkinsonism do not disrupt catalytic activity (67). Moreover, while loss of iPLA_2_-VIA is associated with aberrant mitochondrial morphology in neurons, it also is accompanied by defects in numerous other processes, including Ca^+2^ homeostasis, ER membrane equilibrium, and vesicle trafficking (26–28, 34, 39). Thus, as with other forms of neurodegeneration, further investigation is necessary to determine which defects are drivers and which are passengers in PLAN.

## Materials and Methods

### Drosophila strains

*Drosophila* were raised on standard media at 23°C. *yw (BDSC_6599*)*, P[EPgy2]iPLA2-VIA^EY5103^ (BDSC_15947), Df(3L)BSC394 (BDSC_24418), Df(3L)BSC282 (BDSC_23667), pcyt1^16919^ (BDSC_7319*)*, sws^4^ (BDSC_28121), HMS01544 (BDSC_36129), UAS-mCherry-mitoOMM (BDSC_66532* and *BDSC_66533), Psqh-mito-EYFP (BDSC_7194), DJ667-GAL4 (BDSC_8171),*and *pink1-RNAi (HMS02204, BDSC_41671)* were from Bloomington *Drosophila* Stock Center; *P[GSV6]GS15374* (*DGGR_206187*) was from Kyoto *Drosophila* Genomics and Genetic Resource Center; *tubulin-GAL4* and *elav-GAL4* were gifts from J. Treisman; *bam-GAL4-VP16* was a gift from Y. Yamashita; *NGT40-GAL4* was a gift from R. Lehmann; *eas^KO^* and isogenic controls were gifts from the Jan lab (44); *taz ^161^* mutants were a gift from M. Schlame and M. Ren (68).

The *iPLA_2_-VIA ^23^* mutant was generated by excision of the P element EY5103, which is marked with mini-w+. The location of EY5103 was confirmed by inverse PCR before excision. Over 450 w-lines were screened for lesions in the region of interest by PCR with primers flanking the insertion site (F: 5’-CCGTGCTCTGTGGAATCAGT-3’, R: 5’-GCGCCAAAGCTAGAATTCCG-3’). Mutant line *^Δ23^* was identified in the PCR screen, and the genomic region of *iPLA_2_-VIA* was sequenced to confirm the lesion. Several precise excision lines from the screen were kept as isogenic controls and confirmed by sequencing to be wild-type in the *iPLA_2_-VIA* gene region. This sequencing also confirmed that the same polymorphisms were present in the *^Δ23^* line and the precise excision lines. Revertant line Δ11 was used as the isogenic control in all experiments described here. All recombinant chromosomes made with the *iPLA_2_-VIA ^23^* allele were confirmed by PCR.

### RT-PCR

Whole fly RNA was prepared from mixed sex samples (5-15 flies per sample, flies < 1 week old) with Trizol reagent (Invitrogen) according the manufacturer’s protocol. RNA preparations were treated with DNase I to remove genomic DNA (New England Biolabs) and quantified using NanoDrop (ThermoFisher). cDNAs were reverse transcribed using Superscript III (Invitrogen) according to the manufacturer’s protocol, from an equal mass of RNA for each sample per experiment (1-1.5 µg).

Qualitative PCRs were 25 cycles, unless noted otherwise. Experimental (*iPLA_2_-VIA*) and control primers (*actin*) were included in the same tube to control for pipetting. All experiments were repeated in triplicate. Quantifications were performed using Bio-Rad ImageLab 5.0. PCR primers were as follows:

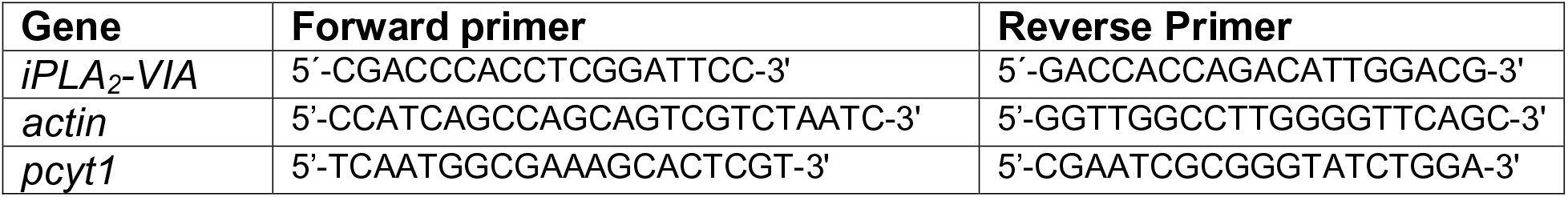

Quantitative PCRs were performed on cDNA samples diluted 6x, using SYBR green + regular ROX mix (MCLab, HSM-400). Reactions were run on an Applied Biosystems 7300 Real-Time PCR System. Primers were:

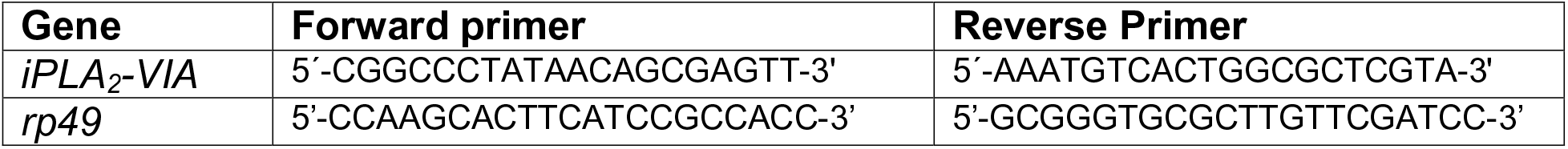

### Climbing assays

Climbing tests were performed as in (69). Groups of 6-13 male or female flies were tapped to the bottom of a fresh food vial and given 20 s to climb 6 cm, into a new empty vial placed on top of the old one. Each group was given five climbing trials per assay. Each fly in the group was given one point for every success, and the total number of points for the group was divided by the number of flies in the group to yield the climbing index. Climbing indices were averaged for at least 6 groups per condition and plotted with standard deviations. Climbing indices for each condition were verified for normal distribution around the average. Statistical comparisons by unpaired t-tests.

### Transgene generation

The wild-type iPLA_2_-VIA-PB transgene was generated by PCR from publicly available iPLA_2_-VIA cDNA (https://dgrc.bio.indiana.edu/, clone RE23733) using Platinum *Pfx* polymerase (Life Technologies, 11708013) and cloned into pUASg-HA.attB (http://www.flyc31.org/) or pPWH (https://dgrc.bio.indiana.edu/vectors/Catalog?product_category=3) using the Gateway system (Life Technologies). PCR primers were: 5’-(CACC)ATGGCGTGGATGGCGTTAG-3’ and 5’-AATCCGTCTTGCATGCGATTTTAG-3’. To introduce the serine-to-alanine mutation, an upstream PCR product was generated with primers 5’-TCTGTACCGGTCGCCGGTG-3’ and 5’-TAGAATTCCGCCAGTAGCGGTGCCGG-3’, and a downstream PCR product was generated with primers 5’-ATTGCCGGCACCGCTACTGGCGGAATT-3’ and 5’-CATGGCGTCTAGAGTCGGGTTGT-3’. The upstream and downstream PCR products were annealed and used as template to generate a PCR product spanning the entire segment. The mutated segment was cloned to replace the corresponding wild-type segment using the unique SgrA1 and Xba1 restriction sites. PCR products were cleaned with QIAquick PCR Purification kit (Qiagen 28104), digested DNA was cleaned with QIAquick Gel Extraction kit (Qiagen 28704), and plasmid DNA was prepared with QIAprep Spin Miniprep kit (Qiagen 27104) and Qiagen Plasmid Midi kit (Qiagen 12143). All constructs were verified by sequencing (Eton Bioscience) before injecting into flies (Genetivision).

### RNA in situ hybridizations

*iPLA_2_-VIA* cDNA plasmid (https://dgrc.bio.indiana.edu/, clone RE23733) was linearized with BamH1 (sense) or Not1 (antisense) restriction digest, and digoxigenin-labeled RNA probes were transcribed using a digoxigenin-UTP in vitro transcription labeling mix (Roche) and either T7 or T3 RNA polymerase (Roche). Transcripts were carbonate-treated to reduce their size.

For imaginal discs, protocol was adapted from (70). Third instar larvae were dissected in PBS and fixed in 4% formaldehyde/PBT (PBT is PBS/0.1% Tween-20) for 1 h at 4°C. Tissues were washed once in 50% PBT:methanol and once in 100% methanol before overnight storage in 100% methanol at −20°C. After two ethanol washes, tissues were incubated in 1:1 xylenes:ethanol for 1 h at 4°C. Tissues were washed again twice in ethanol and once in 50% methanol, and then incubated in 80% acetone for 10 min at −20°C. Tissues were washed twice in PBT and re-fixed in 4% formaldehyde/PBT for 20 min at 4°C. After three PBT washes, tissues were equilibrated into hybridization buffer and hybridized overnight at 65°C with digoxigenin-labeled RNA probes. The next day, tissues were equilibrated back into PBT, washed extensively, and incubated with alkaline phosphatase conjugated anti-digoxigenin antibody (1:1000, Roche) for 1 h at room temperature. After several PBT washes, tissues were washed in AP buffer and color was developed using NBT and BCIP substrates. Hybridization and color development were performed as in (71). Larval and imaginal tissues were mounted in 80% glycerol.

For ovaries, protocol was performed as in (71).

For testes, protocol was performed as in (72). Stained testes were mounted in 60% glycerol:PBS.

### Tissue staining and immunofluorescence

For phalloidin staining, testes were dissected in PBS and fixed in 5% formaldehyde/PBX (PBS/0.1% Triton X-100) for 20 min at room temperature, washed in PBX for 15–20 min, and stained with rhodamine-phalloidin (1:200, Sigma-Aldrich) and DAPI (1:4000, Roche) for 20 min at room temperature. Following three washes in PBX, tissues were mounted in Fluoromount G (Southern Biotechnology).

For antibody staining, testes were fixed and washed as above. Testes were blocked in PBS + 5% normal donkey serum + 1% Triton X-100 before the primary antibody incubation. Primary antibody incubations were performed overnight (in PBX + 5-10% normal donkey serum) at 4°C, washed three times in PBX, and incubated with secondary antibodies, DAPI, and rhodamine-phalloidin for 2 hours at room temperature.

Larvae were dissected in PBS and fixed in 4% formaldehyde for 30 min at 4°C. Tissues were blocked in PBS + 5% normal donkey serum before the primary antibody incubation. For ovary antibody staining, adult females were mated to males on live yeast paste for 1-2 days before dissections. Ovaries were dissected in PBS and combed open, fixed in 5% formaldehyde for 13 min at room temperature, rinsed in PBX, washed 3-4 times in PBX, and blocked in PBS + 5% normal donkey serum + 1% Triton X-100 before primary antibody incubation. Antibody incubations were performed as above.

Primary antibodies used were mouse anti-ATP-5A (1:1000, Abcam), mouse anti-Calnexin 99A (1:2, Developmental Studies Hybridoma Bank, ER marker), mouse anti-Golgin 84 (1:2, Developmental Studies Hybridoma Bank, Golgi marker), rat anti-HA (1:100, Roche 3F10), rabbit anti-cleaved caspase 3 (1:100, Cell Signaling Technology 9664), and rabbit anti-GFP (1:5000, Life Technologies A6455). Secondary antibodies were Alexa Fluor® 488-AffiniPure Donkey Anti-Rat or Anti-Rabbit IgG and Cy3-AffiniPure Donkey Anti-Mouse IgG or Anti-Rat IgG (1:200, Jackson ImmunoResearch).

Images were captured using an Olympus IX-81 motorized inverted microscope with XM-10 monochrome camera (lenses: 10x/0.3 NA, 20x/0.75 NA, 40x oil/1.3 NA, 60x oil/1.35 NA), Zeiss LSM510 Confocal (lenses: 10x/0.3 NA, 20x/0.8 NA, 40x/0.75 NA, 40x oil/1.30 NA), or Zeiss LSM800 Confocal (lenses: 10x/0.3 NA, 20x/0.8 NA, 40x oil/1.3 NA). ICs were scored as described in (73).

### Testis squashes

Internal reproductive organs were dissected from newly eclosed males in PBS, placed in a drop of PBS on a coverslip and punctured. A glass microscope slide was placed on top of the coverslip to squash the testes, and phase-contrast images were collected within 30 min of dissection using an Olympus IX-71 inverted microscope with Nikon Digital Sight camera with DS-U2 controller (20x lens/0.40 NA).

### Fertility tests

For male fertility tests, ten males of each genotype were mated individually to two 3-4 day old *yw* virgin females at 23°C. The males were discarded after one day, and the females were transferred to fresh vials each day for three more days and discarded on day 5. Progeny per vial were counted and averaged.

For homozygous female fertility tests, ten virgin females of each genotype were mated individually to two *yw* males at 23°C. All flies were 3 days old at the time of mating. For hemizygous female fertility tests, virgin females less than one week old were mated to 1-day old *w^1118^* males at 23°C. The males were discarded after one day, and the females were transferred to fresh vials each day for three more days and discarded on day 5. Progeny per vial were counted and averaged.

For egg laying tests, 20 virgin mutant or control females were mated individually to one 7-day old *w^1118^* male each at 23°C. Young females were 7 days old at the time of mating; aged females were 18-19 days old at the time of mating. The males were discarded after one day, and the females were transferred to fresh vials each day for three more days and discarded on day 5. Eggs per vial were counted daily.

Maternal lethality tests were conducted by mating 20-25 young (<10 day old) *iPLA_2_-VIA^Δ23^* or isogenic control females to control males and transferring them to cages with grape juice plates and live yeast paste overnight at 23°C. After 20 hours of egg-laying, 90-120 eggs from each genotype were collected per experiment and moved to new plates at 23°C. Larvae were collected and counted every 24 hours at 23°C. Third instar larvae were moved to vials, and pupae were counted at 23°C. The results of three experiments were averaged.

### Mito-YFP quantification

*iPLA^2^-VIA^Δ23^* mutant (*Psqh-mito-EYFP, iPLA^2^-VIA^Δ23^/iPLA*^2^*-VIA^Δ23^)*, heterozygote (*Psqh-mito-EYFP, iPLA_2_-VIA*^Δ^*/revertant*Δ*11*), and control females (*Psqh-mito-EYFP/revertant*Δ*11*) expressing *Psqh-mito-EYFP* were aged in groups of 10-20 flies at 23°C. Females were mated to males on live yeast paste 1-2 days before dissection. Ovaries were dissected and stained with anti-GFP antibodies as described above. Every stage 8 or 9 egg chamber from each sample was photographed using identical confocal settings and 40x oil objective. After drawing an ROI around the nurse cell area and removing background signal with the same threshold, each image was converted to binary in ImageJ. Each binary image was converted to outlines, and the raw integrated density was measured for the ROI.

### JC-1 fluorimetry

18-30 ovaries per sample were dissected from age-matched *iPLA_2_-VIA^Δ23^* mutant, heterozygote, and control flies (genotypes and aging conditions as above). Tissues were immediately transferred to cold 1x PBS. Sigma mitochondrial isolation kit (Sigma, MITOISO1) was used to extract mitochondria and perform the JC-1 assay according to the manufacturer’s protocol, as follows. After discarding the PBS, each tissue sample was rinsed twice with cooled 1x Extraction Buffer using a glass Pasteur pipet. In a 3 ml glass homogenizer, the lysate was prepared in 1 mL of cooled 1x Extraction Buffer containing 2 mg/ml albumin and transferred to a microcentrifuge tube. The lysate was spun at 1000 x g for 5 minutes at 4°C to remove cell debris, and the transferred supernatant was spun at 13,000 x g for 10 minutes at 4°C to obtain the mitochondrial pellet. The pellet was resuspended in cooled 1x Extraction Buffer (without albumin) by pipetting, and the two centrifugation steps were repeated to get the mitochondria-enriched pellet, which was resuspended in cooled 1x Storage Buffer and stored on ice for immediate use or at 4°C for a maximum of 24 hours. Mitochondrial protein concentration was determined by Bradford assay using a Beckman DU800 spectrophotometer. Sample concentrations were equilibrated with storage buffer. JC-1 assays were performed in clear bottom 96-well plates using a Beckman Coulter DTX800 spectrofluorometer. Each JC-1 reaction mixture of 100 μl was prepared with the same amount of mitochondrial sample (5-7 μg of mitochondrial protein per reaction), 1 μl of the JC-1 solution (final concentration 10 μg/ml) and 0.5 μl of the electron transport chain complex III inhibitor Antimycin A (final concentration 200 μg/ml) or 0.5 μl of the vehicle DMSO. Three technical replicates were prepared for each sample along with a blank lacking mitochondrial sample. After 10 minutes in the dark at room temperature, fluorescence intensity was measured using excitation wavelength of 485 nm, emission wavelength of 595 nm, and integration time of 1000 ms. The average fluorescence per mg of protein of each biological sample was calculated across technical replicates and converted to relative percentage of the isogenic control. For each genotype, 3-4 biological replicates were averaged.

## Acknowledgements

We are grateful to the Bloomington *Drosophila* Stock Center (supported by NIH grant P400D018537), Kyoto *Drosophila* Genomics and Genetic Resource Center, Developmental Studies Hybridoma Bank (created by the NICHD of the NIH and maintained at The University of Iowa, Department of Biology, Iowa City, IA), *Drosophila* Genomics Resource Center (supported by NIH grant 2P40OD010949), Jessica Treisman, Yukiko Yamashita, Ruth Lehmann, the Jan lab, Michael Schlame, and Mindong Ren for fly stocks. We are grateful, as always, to Flybase, for its continual support of the *Drosophila* community (FlyBase is supported by NIH grant U41HG000739 from the National Human Genome Research Institute). Sarah Liberow and Eliezer Heller assisted with some of the experiments. Critical comments on the manuscript were provided by Jessica Treisman and Rebecca Delventhal. Funding for this project provided by NICHD grant R15HD080511 to J.S. and the Yeshiva University Provost’s Office.

