## Supplemental Data for "iPLA_2_-VIA is required for healthy aging in neurons, muscle, and female germline in *Drosophila melanogaster*"

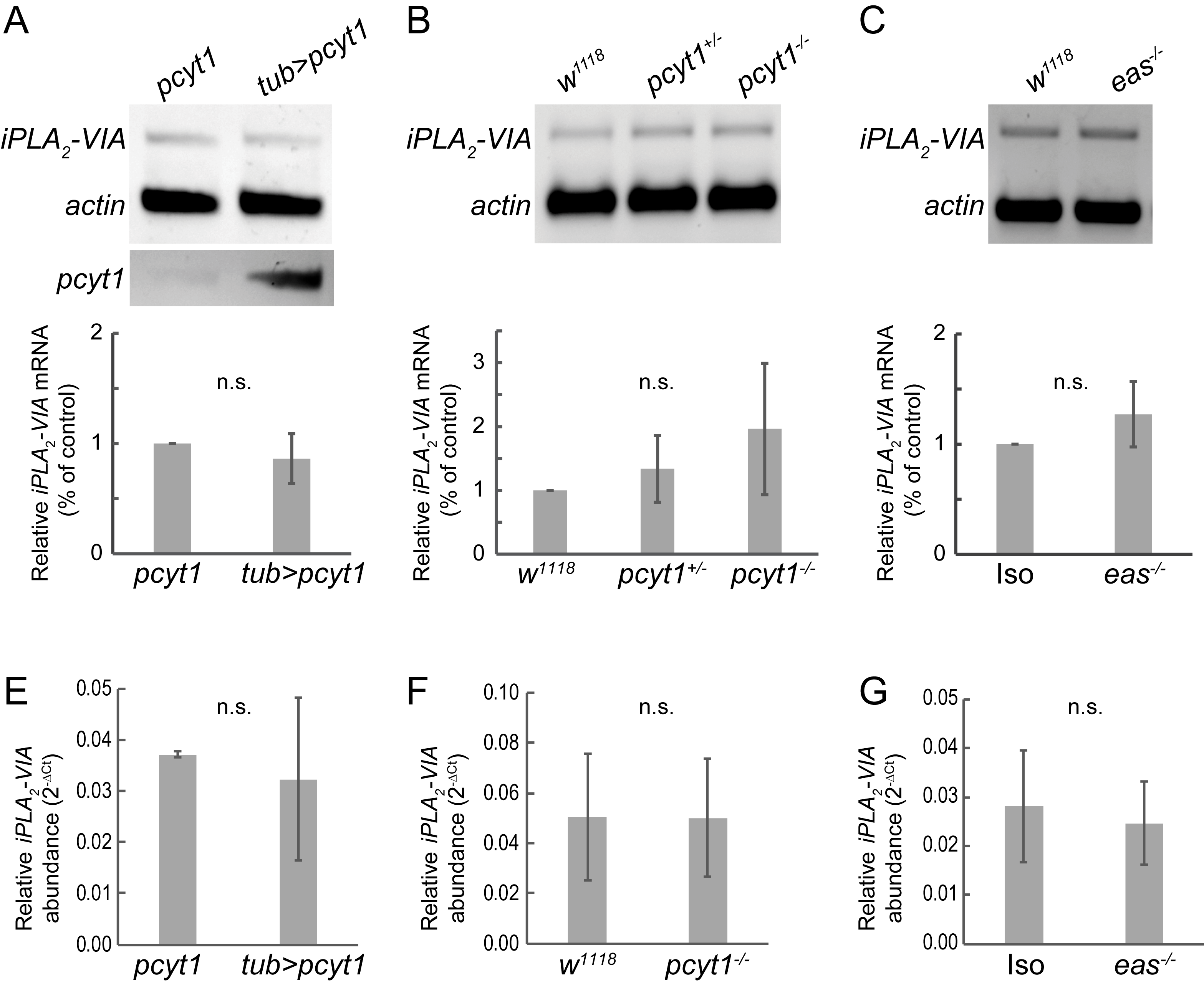

**Figure S1.** ***iPLA_2_-VIA* mRNA levels are unchanged by genetic perturbation of phospholipid metabolism genes.** (A) Whole fly reverse transcription (RT) PCR shows no upregulation of *iPLA_2_-VIA* mRNA expression when Pcyt1 is overexpressed using *tubulin-GAL4*. Control siblings lack the *tubulin-GAL4* driver. *pcyt1* mRNA upregulation is shown in the bottom panel of the gel image. (B-C) Whole fly RT-PCR shows no downregulation of *iPLA_2_-VIA* expression in either the *pcyt1^16919^* mutant compared to *w^1118^* (B) or in the *eas^KO^* mutant compared to isogenic controls (C). Experiments were performed in triplicate. Quantifications were taken with Bio-Rad ImageLab 5.0, shown below representative gel images. Graphs show the ratio of *iPLA_2_-VIA* mRNA normalized to internal control *actin* mRNA in each mutant genotype compared to the control genotype, averaged across three biological replicates. Error bars are standard deviations. (E-G) Whole fly RT-qPCR confirms that *iPLA_2_-VIA* mRNA levels are not significantly different from controls when (E) PCyt1 is overexpressed using *tubulin-GAL4,* (F) in *pcyt1^16919^* mutants, or (G) in *eas^KO^* mutants. *iPLA_2_-VIA* mRNA abundance was normalized to *rp49* mRNA abundance (2^-ΔCt^) and averaged across the three biological replicates. Error bars are standard deviations. Statistical analysis by unpaired t-test (A, C, E-G) or single factor ANOVA (B).

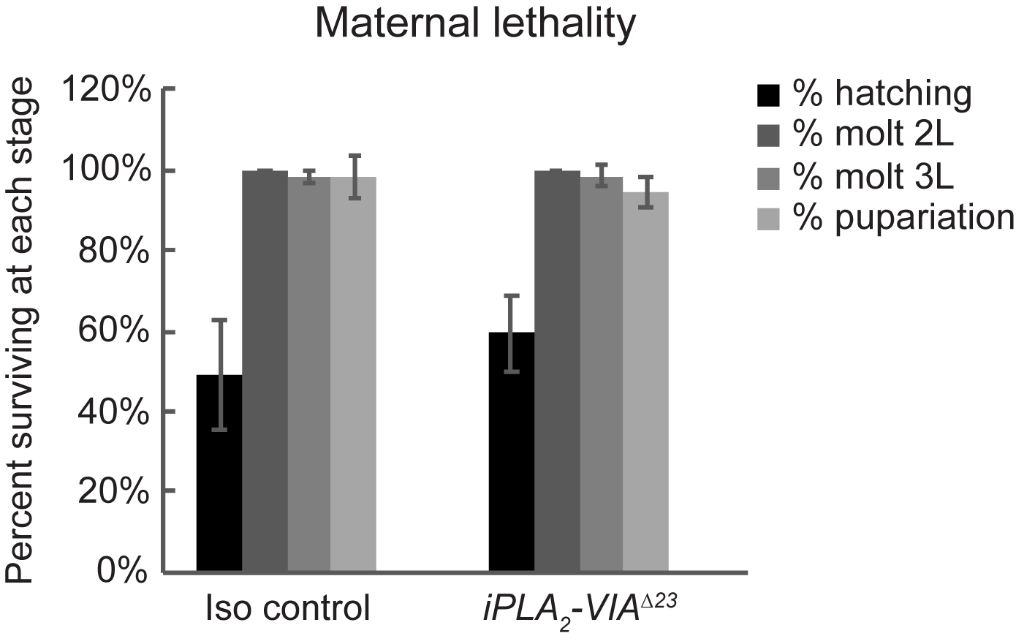

**Figure S2. No maternal effect lethality in *iPLA_2_-VIA^∆23^* mutants.** Young (<10 day old) homozygous *iPLA_2_-VIA^∆23^* females or isogenic control females were mated to control males and allowed to lay eggs on grape juice plates for 20 hours at 23^o^C. First instar larvae hatched from isolated eggs were counted (black bars). Second instar larvae molted from isolated first instars were counted (dark gray bars). Third instar larvae molted from isolated second instars were counted (medium gray bars). Pupae were counted from isolated third instars (light gray bars). The entire experiment was repeated three times. Bars represent the average percentage of individuals that progress to each stage in the three experiments. Error bars are standard deviations. No developmental lethality is observed.

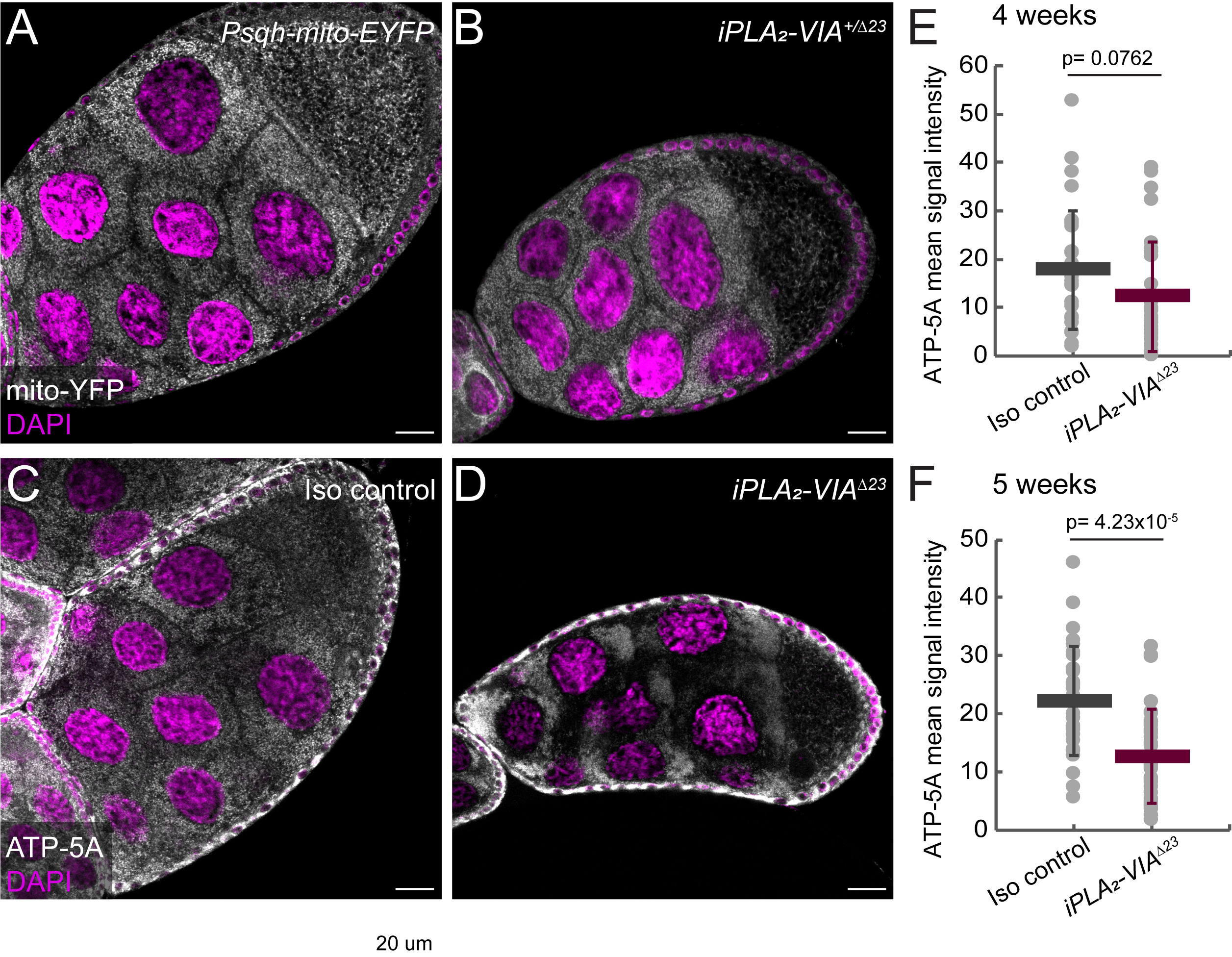

**Figure S3. Mitochondrial aggregation is characteristic of aged *iPLA_2_-VIA* mutant germ cells.** (A-B) Neither the parental stock carrying homozygous *Psqh-mito-EYFP* (A) nor *iPLA_2_-VIA^Δ23^* heterozygotes (B, *Psqh-mito-EYFP, iPLA_2_-VIA^∆23^/revertant∆11*) show mitochondrial clumping even at 6 weeks of age (white, mito-YFP; magenta, DAPI). (C-D) Mitochondrial clumping also is observed with another marker, immunofluorescence to ATP-5A protein in *iPLA_2_-VIA^Δ23^* mutants (D) but not in isogenic controls (C) by four weeks of age (white, anti-ATP-5A; magenta, DAPI). (E-F) Additionally, the ATP-5A signal is weaker in *iPLA_2_-VIA^Δ23^* mutants than in age-matched controls at four (E) and five (F) weeks of age, possibly indicating mitochondrial degradation in the mutant. Scale bars: 20 µm.

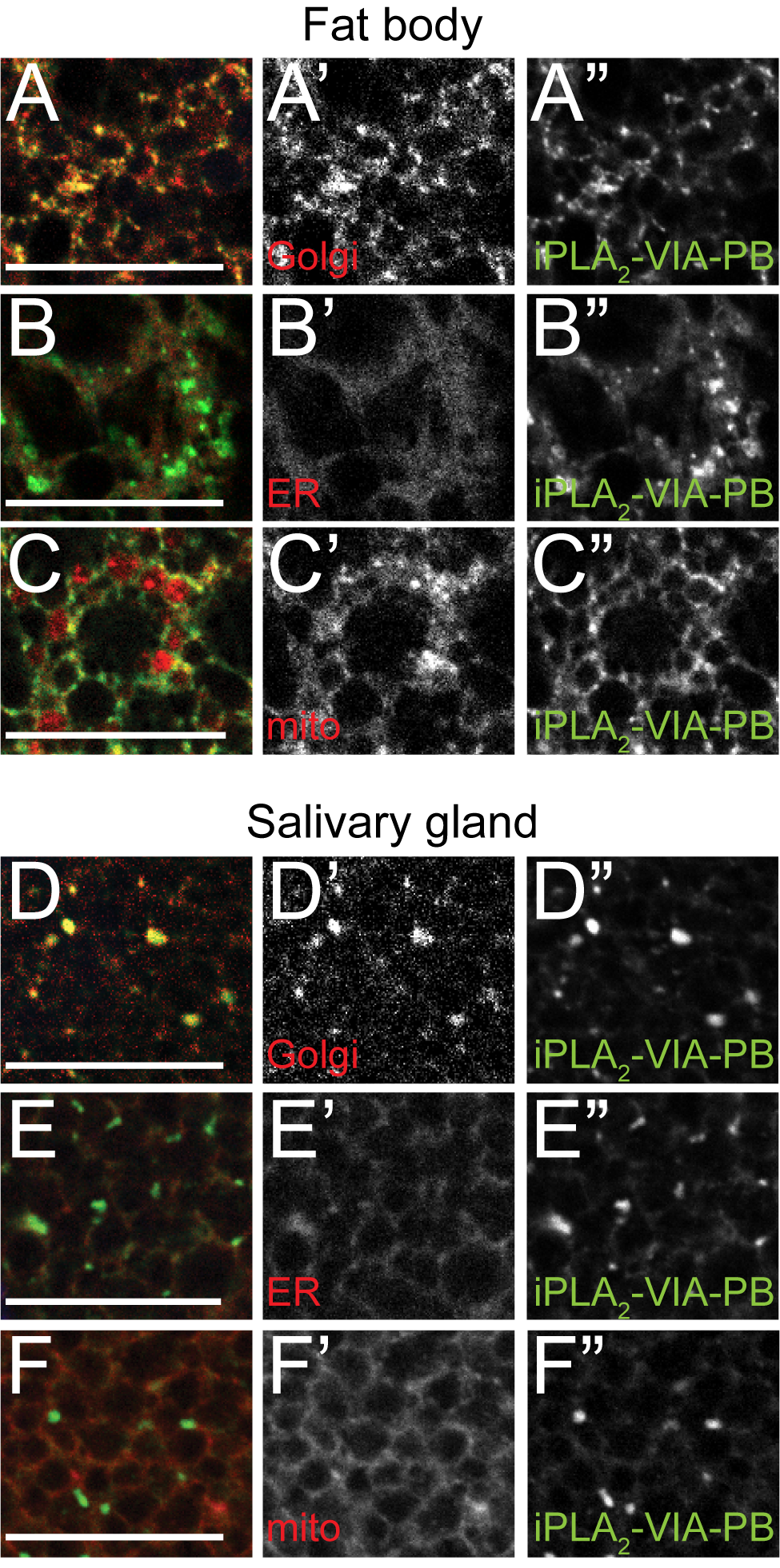

**Figure S4.** **iPLA_2_-VIA-PB localizes to Golgi in somatic larval tissues.** HA-tagged wild-type iPLA­_2_-VIA-PB transgene appears in puncta (green, A”-F”, expressed with *tubulin-GAL4*) that colocalize with a Golgi marker (red, anti-Golgin 84) in larval fat body (A) and salivary glands (D). Colocalization with an ER marker (red, B, E, anti-Calnexin 99A) and a mitochondrial marker (red, C, F, *UAS-mCherry-mitoOMM*) is weak or undetectable in these larval tissues. Scale bars: 20 μm.
